# Single-cell bacterial culturing and antibiotic susceptibility testing using permeable hydrogel-shelled microcapsules

**DOI:** 10.1101/2025.04.03.647129

**Authors:** Shawna L. Pratt, Kerry S. Williamson, Michael J. Franklin, Connie B. Chang

**Affiliations:** Center for Biofilm Engineering, Montana State University, Bozeman, Montana, United States of America; Department of Chemical and Biological Engineering, Montana State University, Bozeman, Montana, United States of America; Department of Microbiology and Immunology, Montana State University, Bozeman, Montana, United States of America; Department of Physiology and Biomedical Engineering, Mayo Clinic, Rochester, Minnesota, United States of America

## Abstract

Microbial communities, such as biofilms, consist of bacteria that exhibit cell-to-cell heterogeneity in their physiological properties, including enzyme activity and gene expression. This single-cell heterogeneity influences the community’s overall metabolic activity and contributes to stress tolerance, such as antimicrobial resistance. To study the impact of single-cell heterogeneity on population-level behaviors, methods have been developed to isolate and characterize bacteria at the single-cell level. One such method includes water-in-oil drop-based microfluidics. However, a limitation of this approach is that the droplet contents cannot be exchanged during experiments, making it difficult to investigate how cells respond to changing environmental conditions. To address this limitation, we developed a drop-based microfluidic technique called Bioflex, which creates permeable hydrogel-shell microcapsules that act as growth chambers for individual bacterial cells. The microcapsules are formed by crosslinking a shell made from a PEG-based hydrogel (4-arm PEG-maleimide) around a dextran core. After cross-linking, the dextran core diffuses out of the capsules and is replaced with buffer or growth medium. By adjusting fluid flow rates during the process, we can control the size and shell thickness of the microcapsules, allowing the production of different capsule architectures. The Bioflex capsules are biocompatible, supporting the encapsulation and growth of *Pseudomonas aeruginosa* by allowing nutrient transport across the capsule shell. Moreover, the capsules remained permeable to antimicrobial treatments introduced during *P. aeruginosa* incubation. For example, *P. aeruginosa* cells in Bioflex capsules responded to externally delivered ciprofloxacin treatments, with their responses varying depending on the timing of the antibiotic introduction. In summary, Bioflex capsules provide a novel, high-throughput platform for isolating single-cells and testing how single-cell derived communities react to time-dependent stresses, such as antibiotic treatments.

**Graphical Abstract:** 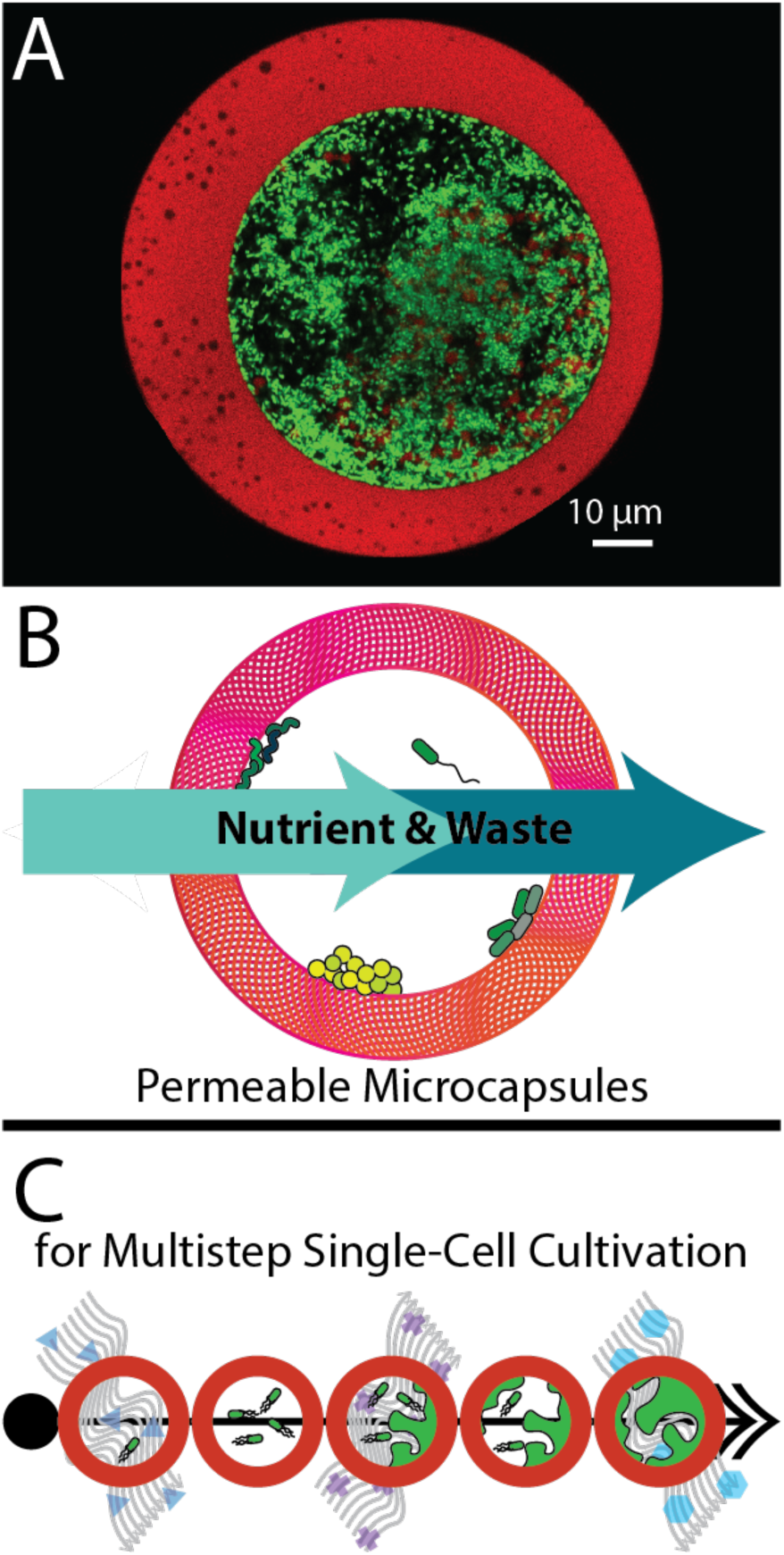

Bacterial cultivation using permeable microscale Bioflex capsules. (A) *P. aeruginosa* PAO1 expressing eGFP cultured in a Bioflex capsule starting from a single cell. (B) Bioflex capsules are permeable to nutrient and waste transport. (C) The capsules allow for exchange of media to study the responses of single cells or small populations of cells within the drops using time-lapse imaging.

## Introduction

Biofilms, defined as microbial communities bound by extracellular matrices, consist of physiologically heterogeneous assemblages of cells. Heterogeneity in microbial communities can lead to antimicrobial resistance and recalcitrance, genetic divergence, or cooperative behaviors^1,2^. In the context of medical microbial infections, these heterogeneity-derived traits can complicate and hinder treatment^3^. Multiple approaches have been developed to characterize bacteria and other microorganisms with single-cell resolution, thereby increasing understanding of heterogeneity within microbial communities. Examples of these approaches include labeling individual cells based on genotype using fluorescence *in situ* hybridization (FISH)^4,5^ or based on cell-level metabolic activity via bioorthogonal noncanonical amino acid tagging (BONCAT)^6^.

Single cells can also be characterized using fluorescent labels in conjunction with fluorescence-activated cell sorting (FACS)^7,8^ or via single-cell DNA or RNA sequencing to determine genetic diversity and expression patterns^9^. Notably, these methods are primarily used for endpoint analysis and are less amenable to tracking viable single-cells or single-cell lineages at multiple time points as cells grow or diversify. Techniques such as microfluidic flow cell imaging^10,11^, compartmentalization^12^, and microenvironment assays^13^, provide single-cell resolved imaging data.

Drop-based microfluidic approaches^14–17^ are uniquely valuable for high-throughput sample generation and single-cell resolution in longitudinal time-resolved studies. Single cells in various media conditions can be isolated inside water-in-oil drops^14,15^. These drops can then be collected and incubated in a microfluidic chamber for time-lapse imaging to subsequently characterize heterogeneity in growth kinetics^14,15,17^. However, cultivation in water-in-oil drops presents two distinct drawbacks: (1) the drop storage environment must be carefully managed to maintain drop stability for time-lapse studies^15^, and (2) the water-in-oil drop interfaces have reduced permeability to water-soluble nutrients and waste. As a result, microbial growth in drops is impacted by nutrient depletion and the accumulation of bacterial waste^16^.

Additionally, due to limitations in molecular transport across water-in-oil interfaces, altering the composition of drops during an experiment to introduce a new nutrient condition or antimicrobial environment may require complex droplet processing steps such as picoinjection or droplet merging^18,19^.

A chemically permeable, single-cell isolation and cultivation platform would enable studies on single bacterial cells, where the chemical environment surrounding the cells could be modified over time as an intentional experimental variable. Hydrogel-shelled capsules are particularly appealing for microbial single-cell cultivation due to (1) the permeability of hydrogels to aqueous media and (2) the ability to isolate planktonic single cells within a hydrogel shell without imparting mechanical stress from embedding in a hydrogel network. One study demonstrated bacterial growth in polyethylene glycol– diacrylate (PEG-DA) capsules^20^. This approach used diacrylate modified PEG, which relies on photo-illumination and free-radical crosslinking chemistry, and a non-coaxially flowing device with crosslinking agents included in the aqueous phase, which requires the precise tuning of pH and polymer concentrations to form a core-shell structure. A similar non-coaxially flowing microfluidic device and crosslinker delivery approach is adapted for use with PEG-maleimide to culture algae cells^21^. PEG-maleimide is preferred over other PEG derivatives for cell and tissue cultivation due to the biocompatibility of the crosslinking process^22^, which proceeds at physiological pH, produces no by-products, and does not rely on free-radical based crosslinking cascades ^23,24^. Finally, a separate approach that does not use phase separation is used to generate PEG-maleimide and PEG capsules using a three-dimensional (3D) coaxially flowing device with crosslinking agent delivered externally to the aqueous fluid streams through a nano emulsion^25^.

In this study, we present a 3D microfluidic device that geometrically templates an inner core dextran phase and an outer shell PEG-maleimide phase solution for rapid crosslinking with an external DTT nanoemulsion. The microfluidic system enables the formation of hydrogel-shelled microcapsules with highly tunable sizes and shell thicknesses. These microcapsules, termed Bioflex capsules, are permeable to aqueous media, biocompatible, and optimized for single-cell microbial isolation and cultivation.

By integrating polymer phase separation with 3D device geometry for efficient chemical crosslinking, we demonstrate highly controllable and tunable hydrogel-shell microcapsule formation. combining the advantages of the hydrogel microcapsule generation methods referenced above^20,21,25^. Shell thickness and capsule size can be precisely adjusted during capsule production without requiring modifications to the device design or capsule precursor formulation. This flexibility enables a longitudinal, single-cell resolved microbial cultivation approach, allowing the use of multi-step processes or transient chemical conditions as time-varied parameters. To validate these capabilities, we evaluated the growth of *Pseudomonas aeruginosa* within the capsules. Additionally, using a multi-step experimental design, we observed how capsule-isolated single-cell lineages respond to variations in antibiotic treatment schedules. These findings highlight the versatility of the Bioflex method for generating highly tunable capsules, isolating and cultivating bacteria, and analyzing microbial responses to changes in environmental conditions.

## Methods

### Core-Shell Templating Device Description and Fabrication

A microfluidic device was developed to generate Bioflex microcapsules, featuring channel and junction architectures modified from previously reported hydrogel-capsule generation devices ^25,26^. To improve performance, channel widths and lengths were adjusted to reduce backflow of crosslinking solutions into the hydrogel precursor channels. Specifically, the core channel was lengthened through serpentine channels and narrowed relative to other channels, increasing resistance, while oil channels were widened and shortened to reduce resistance (Supplementary Figure 1). Computer-aided design and fluid flow analysis were used to model and visualize the device using Autodesk Inventor, AutoCAD, and COMSOL (Supplementary Figure 1).

The Bioflex capsule generation devices were fabricated following standard approaches for PDMS-based non-planar flow-focusing device fabrication approaches^27^. Device masters were created by patterning SU-8 3050 photoresist (Kayaku Advanced Materials) onto 4-inch silicon wafers (UniversityWafer, Inc.) using mylar patterning masks (ArtNetPro) and a photolithography contact aligner (ABM). Separate masters were produced for the top and bottom halves of the non-planar junction. For the top master, core channels were patterned in a 45 µm thick SU-8 layer. Then, an additional 10 µm thick layer of SU-8 was added to the substrate, followed by patterning of the non-core channels in the resultant 55 µm thick layer of SU-8. The completed master was developed thereafter. The bottom master consisted of a single 10 µm thick layer of photoresist. The thickness of each layer and x-y feature fidelity of the masters were verified using optical profilometry (Filmetrics ProFilm 3D). The architecture of both masters is shown in Supplementary Figure 1.

The top and bottom master molds were used together to fabricate slabs of Sylgard 184 polydimethylsiloxane (PDMS, DOW), enabling the production of 14 microfluidic devices simultaneously. The bottom PDMS slab was kept thin (1-3 mm) to ensure compatibility with the working distances of the inverted, non-immersion inverted microscope objectives (4x - 20x). After curing, the PDMS slabs were cut from the silicon wafer masters. Ports were punched in the top slab using a 0.75 mm biopsy punch (Electron Microscopy Sciences EMS-Core Sampling Tool).

The final devices were assembled through plasma bonding of the PDMS slabs to glass slides. For bonding, individual devices were cut from the PDMS slabs corresponding to the top and bottom portions of the microfluidic device. Each slab consisted of a smooth face and a featured face containing depressions that formed the device channels. These PDMS slabs, along with a 1 x 3 inch glass slide (VWR) was exposed to oxygen plasma (Harrick). To ensure plasma reached both the top and bottom surfaces of the bottom PDMS slab, the slabs were balanced on a tinfoil shelf.

After plasma treatment, the smooth side of the bottom slab was bonded to the glass slide. A drop of 0.2 µm filtered distilled water was applied to the featured face of the bottom slab. The top slab was then placed on the bottom face, with a thin film of water facilitating alignment. Without this water layer, the top and bottom slabs would irreversibly bond upon contact. The features of both slabs were visualized using brightfield microscopy and aligned by hand.

After alignment, the devices were cured at 65°C for a minimum of 12 h. The internal channel surfaces of the finished devices were rendered hydrophobic by filling the channels with a 0.2 µm filtered solution of 10 µL/mL (tridecafluoro-1,1,2,2-tetrahydrooctyl) trichlorosilane (Gelest) in Novec HFE 7500 fluorinated oil (3M). The filled devices were baked for 1 h at 65°C, during which the fluorinated oil completely evaporated. For flow characterization via inverted confocal laser-scanning microscopy (iCLSM; Figure 1C), the devices were bonded to a 25 x 40 mm No. 1.5 cover glass (Electron Microscopy Sciences) instead of a 1 x 3 inch slide.

**Figure 1:**
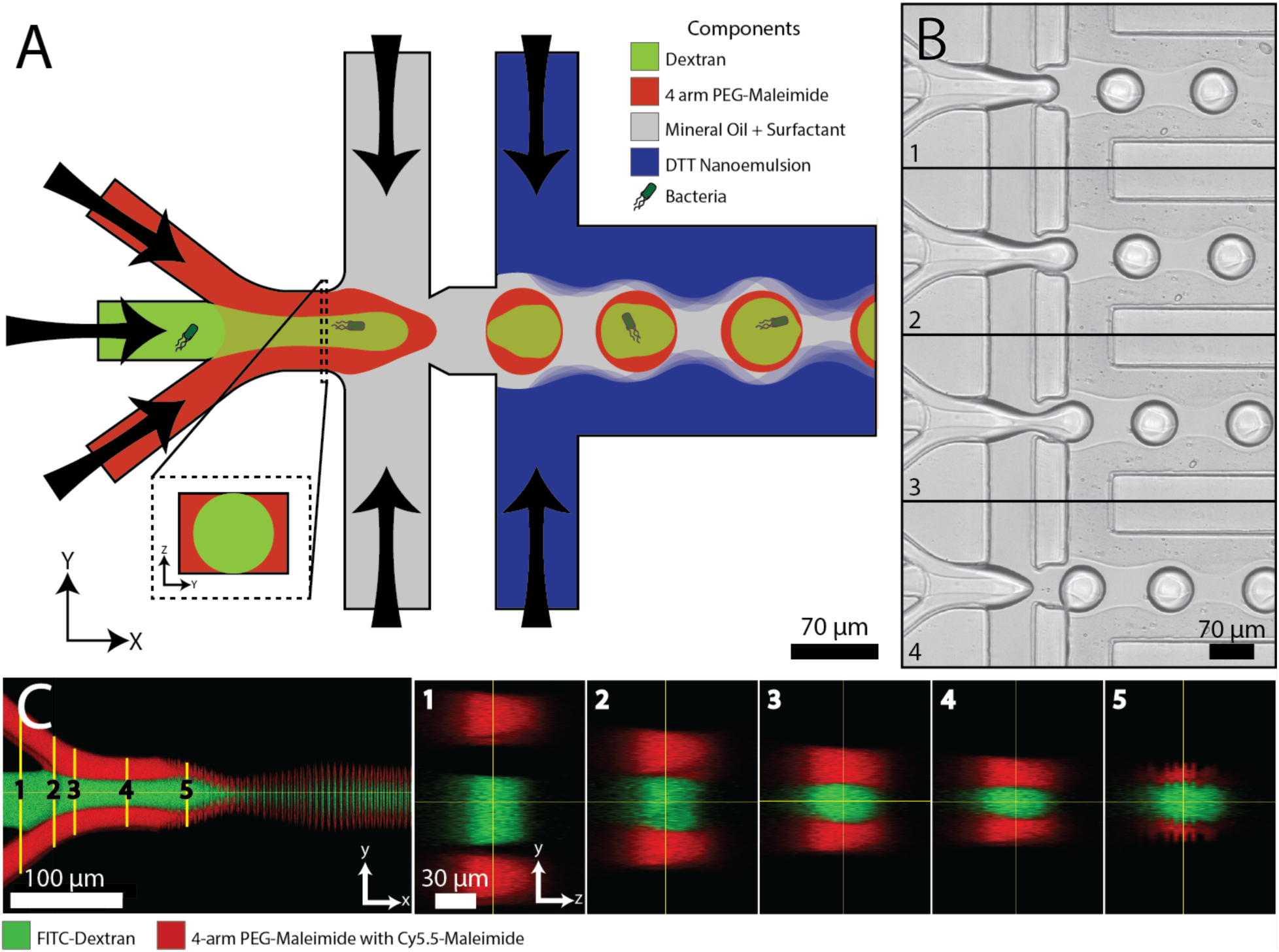
The Bioflex method utilizes drop-based microfluidics for high-throughput generation of hydrogel-shelled, bacteria-encapsulating capsules. (A) Capsules are generated in a co-axial, non-planar flow-focusing microfluidic device using four fluids: aqueous dextran (green), 4-arm PEG-mal (red), mineral oil/surfactant solution (gray), and DTT nanoemulsion (blue). Bacteria are incorporated into the dextran solution for encapsulation. The device is designed to establish a core-sheath upstream of droplet breakup (inset). (B) High-speed imaging captures the droplet breakup sequence, with frames 1 through 4 taken at approximately 1 ms intervals. (C) Orthogonal views of dextran (FITC-labeled) and 4-arm PEG-mal (Cy5.5-maleimide-labeled) flowing through the capsule generation device. Optical sectioning images reveal the evolution of core-sheath flow profiles originating from the non-planar, coaxially aligned device design.

### Preparation of Microcapsule Components

All components were sterilized using a 0.2 µm PTFE (CELLTREAT Scientific Products) filter prior to use, unless specified otherwise. The hydrogel precursor solution consisted of 200 mg/mL 10 kDa 4-arm polyethylene glycol-maleimide (4-arm PEG-mal) (Laysan Bio), 20 µL/mL 5.54 mg/mL Cy5.5-maleimide (Click Chemistry Tools) in DMSO (Sigma-Aldrich), and 1x phosphate buffered solution (PBS) (Becton, Dickinson and Company), pH 7.4. The core solution contained 260 mg/mL 10 kDa dextran (Chem-Impex, Intl.) suspended in 1x PBS for both cell-containing and abiotic studies. For flow dynamics studies during capsule production using iCLSM, FITC-Dextran (10 kDa, 1 wt%) (Sigma Aldrich) was added to the dextran suspension (Figure 1C).

The oil solution was comprised of 3 wt% Span^TM^ 80 (TCI Chemical) dissolved in light mineral oil (Spectrum Chemical). The crosslinking nanoemulsion was prepared by mixing 1-part aqueous solution with 15-parts oil solution. The aqueous solution consisted of 0.2 µm-filtered 50 mg/mL dithiothreitol (DTT) (Thermo Scientific) in 0.2 M triethanolamine (TEA) buffer at pH 8.0 (Thermo Scientific™). The oil solution contained 0.2 µm-filtered 3 wt% Span^TM^80 in light mineral oil solution. To form the nanoemulsion, the aqueous DTT solution and the oil solution were combined in a microcentrifuge tube and emulsified using a wand sonicator. The microcentrifuge tube was kept in a -20 °C microcentrifuge tube cooler during sonication.

### Microcapsule Generation

The flow of microcapsule components (core suspension, hydrogel precursor solution, oil solution, and crosslinking nanoemulsion) was driven from 1 mL or 3 mL Luer-Lok syringes (BD) fitted with 27-gauge Luer-Lok needles (Excel Intl.) through microtubing (PE/2, Scientific Commodities) using syringe pumps (Harvard Apparatus) into the microfluidic device via its four inlet ports, as previously described. The flow rates used in the capsule size, shell thickness, and biological growth studies (Figures 3-6) are listed in Supplementary Tables 1-3. Capsules were collected in a 1.5 mL centrifuge tube (Eppendorf) for 20 min in all biological studies and for 10-20 minutes in abiotic studies. Time-resolved capsule generation (Figures 1B and 3A,D) was visualized using brightfield microscopy on an inverted microscope (Eclipse Ti2 Series Inverted Microscope, Nikon) equipped with a high-speed camera (Phantom).

Following collection, the capsules were washed to replace surrounding mineral oil with an aqueous media and to facilitate the diffusion of dextran from the capsule cores. The washing was as follows: (1) The collection tube was centrifuged for 1 min at 500 x *g*, and the mineral oil supernatant was removed. (2) Fresh mineral oil (1 mL) was added to the tube, vortexed for 15 s or longer to resuspend the capsule pellet, and then centrifuged for 1 min at 500 x *g*. The mineral oil supernatant was removed. Step 2 was repeated twice. (3) Sterile PBS (1 mL) was added to the collection tube, vortexed for 15 s or longer to ensure capsule resuspension, and centrifuged for 3 min at 500 x *g*. After centrifugation, the aqueous supernatant and any residual mineral oil were removed. Step 3 was repeated three times. (4) Finally, 50 µL of PBS was added to the collection tube and vortexed for 30 s to break up any capsule clumps formed during centrifugation.

### Capsule Shell Thickness and Radius Analysis

Capsule shell thickness and radius (Figure 3B, E) were determined from optically sectioned images of the capsules captured via iCLSM (Stellaris DMI-8, Leica). The analyzed optical section corresponded to the mid-plane of the capsules’ 3-D spherical structure. Images were thresholded by pixel intensity value to separate the core, shell, and background into three regions, and the area of each region was quantified. To determine the percent capsule radius, the shell area was divided by the sum of the shell and core areas (Figure 3B). For each sample, 6-8 images were analyzed. Shell thickness was analyzed using base Fiji-ImageJ functionality^28^ . Capsule radius was determined by performing a Hough-transform on capsule images using the UCB vision sciences recommended workflow^29^. Before applying the Hough-transform, images were intensity thresholded to distinguish the shell and core from background. A minimum of 460 capsules per sample were analyzed to determine radii. Statistical analysis and data representation were conducted in Rstudio. Average and standard deviation values for shell thickness and capsule radius were calculated using standard statistical methods.

### Aqueous Two-Phase System (ATPS) Drop Production and Confocal Flow Visualization

iCLSM was used to visualize the formation of water-in-oil drops from capsule precursor solutions (Figure 1C). This allowed for the determination of phase orientation during drop production (Figure 1C) and the equilibrium particle morphology of uncrosslinked mixtures of 4-arm PEG-mal and dextran in drops (Figure 2B.) Aqueous solutions of 10 kDa dextran and 10 kDa 4-arm PEG-mal solutions were prepared in a range of concentrations. Dextran concentrations ranged from 150-300 mg/mL, while 4-arm PEG-mal concentrations ranged from 50-240 mg/mL. The dextran and 4-arm PEG-mal solutions were fluorescently labeled using 1 wt/wt% 10 kDa FITC-dextran (Sigma Aldrich) and 10 µL/mL of 5.54 mg/mL Cy5.5 maleimide (Click Chemistry Tools) in DMSO (Thermo Scientific), respectively.

**Figure 2:**
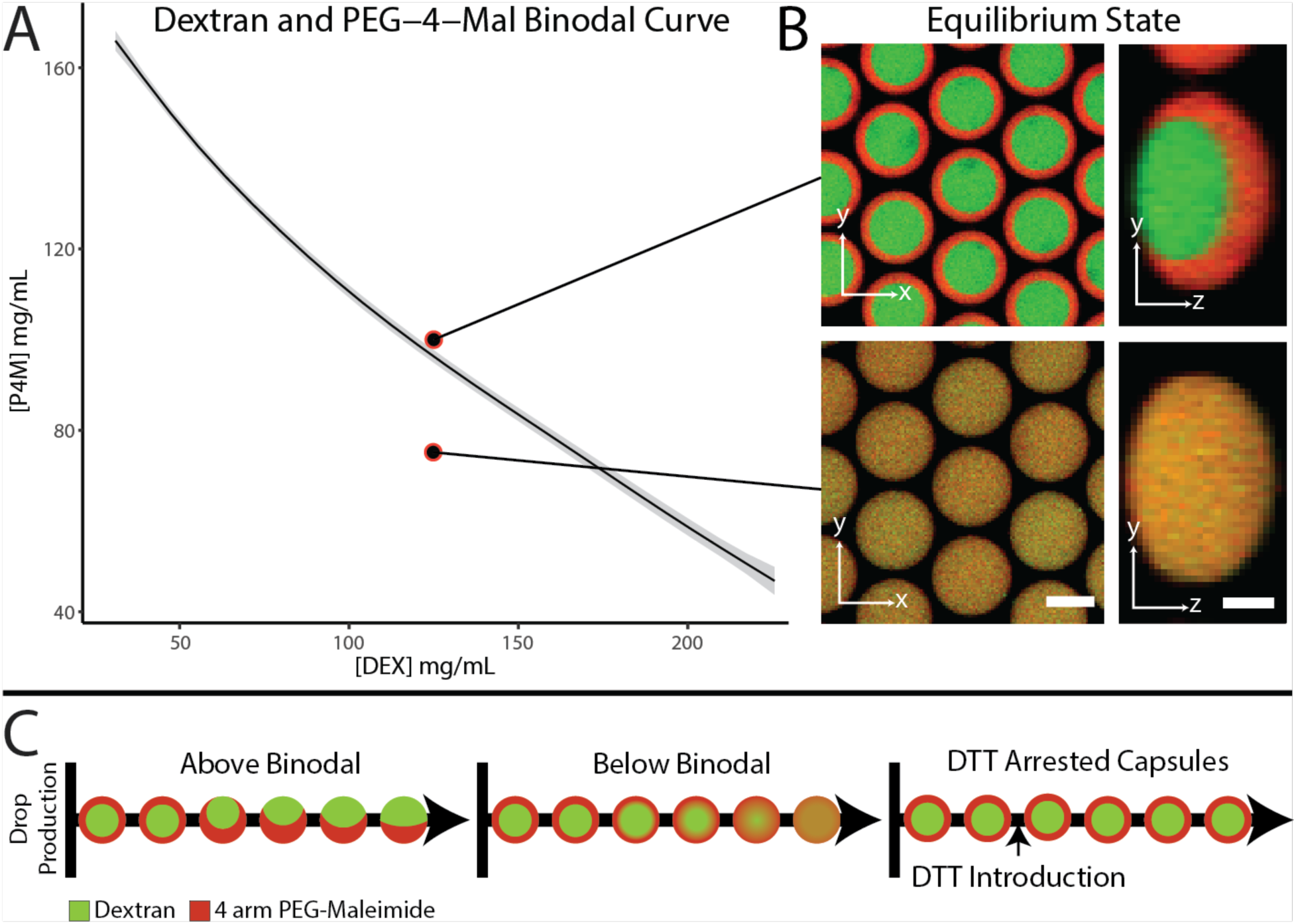
The equilibrium architecture of aqueous drops containing dextran and 4-arm PEG-mal is determined by their position on the phase separation curve. Introducing a crosslinking agent arrests the drop architecture, preventing transition to equilibrium geometry. (A) The binodal phase separation curve for 10 kDa 4-arm PEG-mal and 10 kDa dextran. Mixtures with concentrations below the curve remain homogenous, while those above undergo phase separation. The binodal curve (black line) was determined via cloud point titration (three trials, Loess fit), with gray regions indicating a 95% confidence interval of the Loess fit average. (B) Orthogonal optically sectioned microscopy of uncrosslinked dextran and 4-arm PEG-mal drops with phase separating (top) and miscible (bottom) concentrations of dextran and 4-arm PEG-mal. Top: 125 mg/mL dextran / 100 mg/mL 4-arm PEG-mal. Bottom: 125 mg/mL dextran / 75 mg/mL 4-arm PEG-mal. (C) Schematic illustrating the evolution of 4-arm PEG-mal and dextran drop mixtures in a non-planar microfluidic device, both above and below the binodal curve. Crosslinking arrests phase evolution, stabilizing the core-shell geometry regardless of the drop’s initial position on the binodal curve.

**Figure 3:**
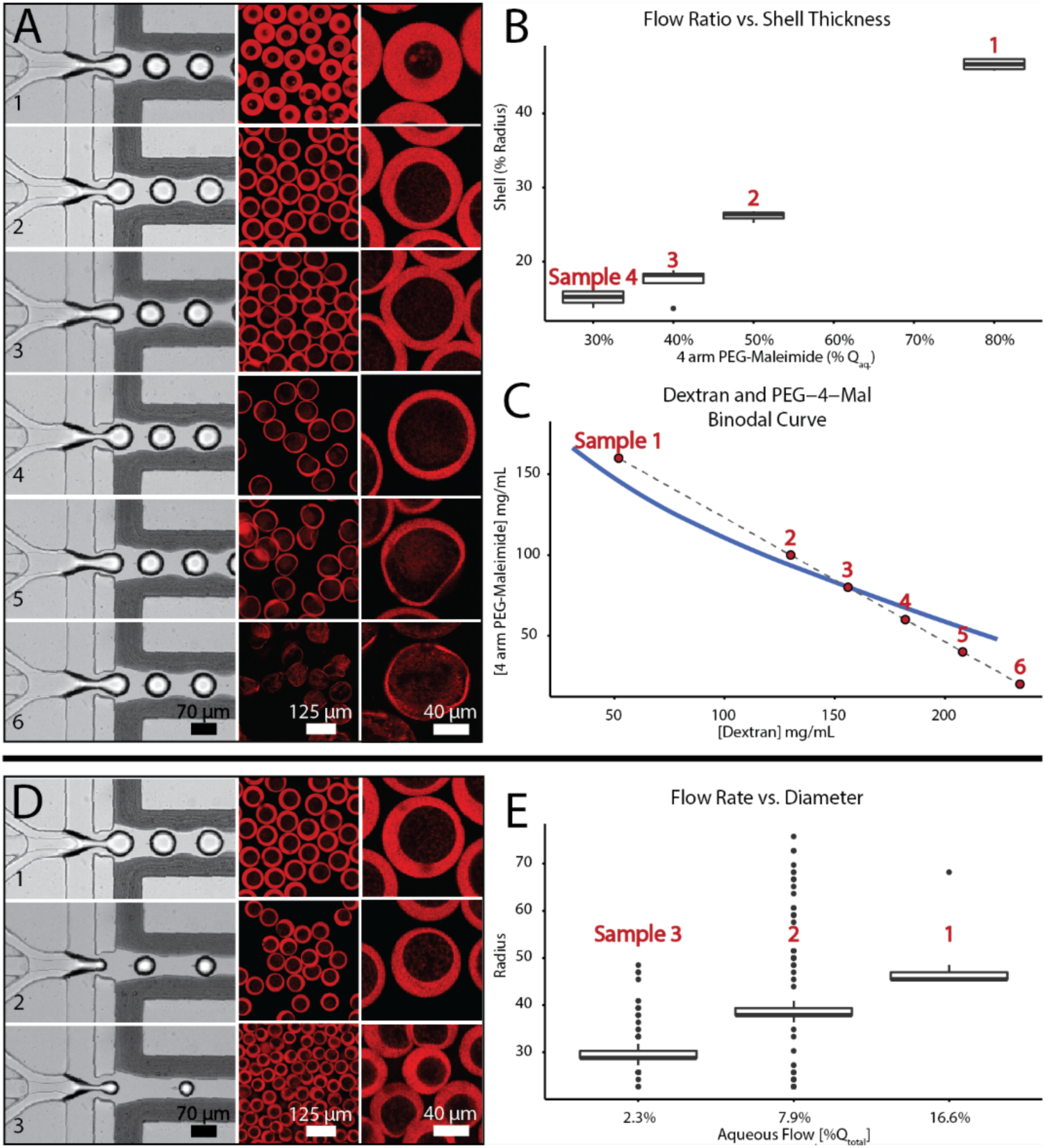
Flow rate controls shell thickness and capsule radius. (A-B) Adjusting the ratio of dextran to 4-arm PEG-mal volumetric flow rates alters capsule shell thickness. (A) The left panels in each series show high-speed camera images of capsule generation. The right panels display iCLSM images taken at the mid-plane of the capsules. Cy5.5-maleimide (red) labels the 4-arm PEG-mal, highlighting the capsule shells. Panels 1-6 correspond to 4-arm PEG-mal solutions making up 80%, 50%, 40%, 30%, 20%, and 10% of the aqueous volumetric flow (*Q_aq_*), with panels 1-3 showing intact capsules. (B) Shell thickness as a function of the dextran to 4-arm PEG-mal flow rate ratio. The y-axis represents the percentage of capsule radius occupied by the shell, while the x-axis represents the percentage of 4-arm PEG-mal solution in *Q_aq_*. (C) The capsule composition in (A) is plotted on the binodal phase separation curve. (D) Modulating the ratio of total aqueous flow (the combined dextran and 4-arm PEG-mal flow rates) to oil flow affects capsule size. The left panels in each series show high-speed camera images of capsule generation. The right panels present iCSLM images of capsules at the mid-plane. 4-arm PEG-mal capsule shells are labeled with Cy5.5-maleimide (red). The aqueous phase constituted 16.6%, 7.9%, and 2.3% of the total volumetric flow in panels 1-3, respectively. (E) Distributions of capsule radii as a function of *Q_aq_* percentage relative to total flow (*Q_total_*).

For both the drop production imaging study and the equilibrium morphology study, drops of 4-arm PEG-mal and dextran were generated using the protocol described in the "Microcapsule Generation" methods section, with one key modification:

the drops were not crosslinked, and therefore DTT nanoemulsion was excluded from the capsule generation process. To accommodate this change, the DTT nanoemulsion inlet on the microfluidic capsule generation device was plugged with a short section of tubing with a sealed end. During drop imaging, 200 mg/mL 4-arm PEG-mal and 200 mg/mL dextran solutions were used. In the equilibrium morphology study, various polymer concentrations were tested. Phase orientation within the droplets was visualized using iCLSM.

### Binodal Phase Separation Studies

The binodal phase separation line between dextran and 4-arm PEG-mail solutions was determined using a cloud-point titration method ^30^ , visually assessed by eye, at laboratory room temperature (19-23°C). No additional temperature control was used to manage fluctuations during the titration and observation process. The titration was performed in three technical replicates (Supplemental Figure 2). To conduct the titration, 170-180 µL of 260 mg/mL 10 kDa dextran in water was dispensed into a spectrophotometric cuvette. 4-arm PEG-mal in water (350 mg/mL) was then added in 5-10 µL increments, with thorough mixing via pipetting. After each addition, the cuvette was backlit with diffuse white LED light against a dark surface, and turbidity was visually assessed. Once turbidity was observed, water was added instead of 4-arm PEG-mal until the solution became clear. This cycle of adding 4-arm PEG-mal followed by water was repeated until the dextran concentration in the cuvette was diluted to below 50 mg/mL. Throughout the titration process, the volumes of dextran solution and water added as well as the cloud and clarity transition points were recorded. Data processing and representation for the binodal diagram were conducted in Rstudio. An average phase separation curve and a 95% confidence interval were determined from the three trials using Loess averaging (Figure 2A).

### Preparation of Bacteria for Encapsulation

Overnight cultures of *Pseudomonas aeruginosa* PAO1 pMF230 ^31^ were prepared by incubating frozen stocks in 125 mL baffled culture flasks containing 20 mL of tryptic soy broth (TSB) (Becton, Dickinson and Company) supplemented with 150 µg/mL carbenicillin (Fisher Scientific). Cultures were incubated at 37 °C with shaking at 160 rpm for 12-15 h. To prepare working cultures, 100 µL of the overnight culture was inoculated into 10 mL of TSB amended with 150 µg/mL carbenicillin in a 125 mL baffled culture flask and incubated at 37 °C with 160 rpm shaking for 4 hours. One mL of the working culture was washed by centrifugation at 5000 × g for 3 min, followed by supernatant removal and pellet resuspension in PBS, pH 7.4 (Gibco, Thermo Fisher Scientific) by vortexing a minimum of 15 s. This process was repeated for a total of three centrifuge cycles. After the final wash, the culture was incubated at room temperature for 30 min to allow for the completion of any ongoing cell division processes.

To prepare the core solution, the washed working culture was diluted 10-fold in PBS. The diluted culture was then added to a dextran-in-PBS solution at a ratio of 25 µL culture in 1 mL, yielding a final dextran concentration of 260 mg/mL in the core solution. The core solution was then used for capsule generation, as described in the "Microcapsule Generation" methods section.

### Imaging and Quantification of Cell Growth

Time-lapse cultivation studies were conducted to evaluate bacterial growth within capsules and to assess the effects of varying the timing of antibiotic introduction (Figures 4-6, Supplementary Figures 3, 5, and 6). For the growth study without antibiotic treatment (Figure 4, Supplementary Figure 3), 10 µL of capsules and 200 µL of undiluted TSB were dispensed into wells of an 8 chambered #1.5 coverglass (Cellvis).

**Figure 4:**
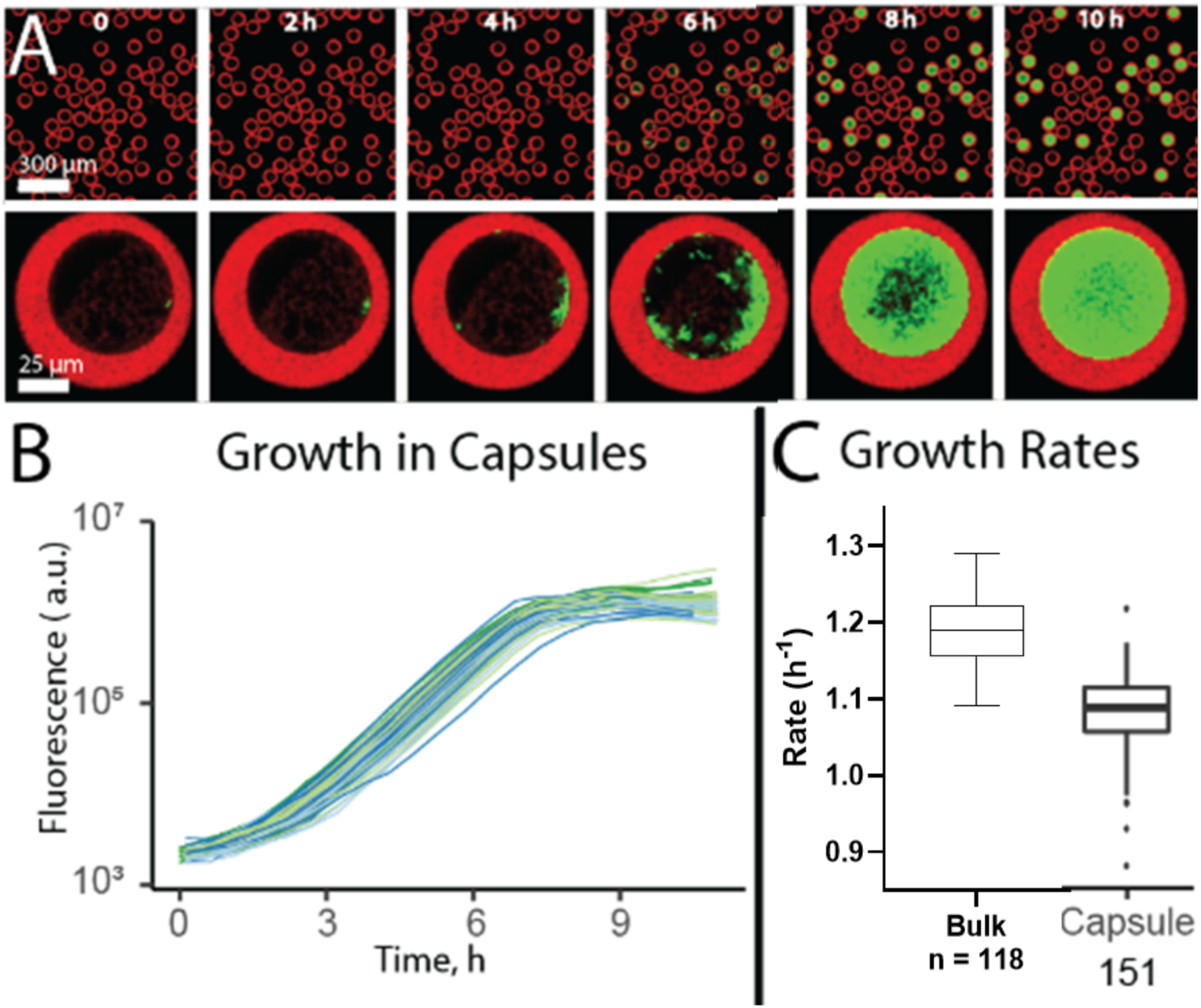
Growth of *P. aeruginosa* PAO1 (pMF230) in Bioflex capsules. (A) Bacterial growth in capsules was monitored over a 10 h incubation period using iCLSM. Capsule shells were labeled with Cy5.5-maleimide (red), while *P. aeruginosa* expressed GFP (green). The bottom panel shows a single capsule that initially contained one bacterial cell, incubated in TSB for 10 h. (B) Growth curves for individual bacteria within capsules were generated based on fluorescence signals. Fluorescence measurements were obtained from iCLSM z-stacks, with fluorescence summed across multiple slices. The plot represents growth curves from a single representative trial; three trials were conducted. (C) Growth rates of *P. aeruginosa* were determined by comparing the doubling time of fluorescence signals in microtiter plate wells and capsules. Sample sizes: *n =* 118 wells across 3 independent biological replicates for microtiter plate assays and *n =* 151 bacteria from 3 independent biological replicates for capsule assays.

For the study examining the effects of antibiotic timing (Figures 5, 6, Supplementary Figures 5,6), 20 µL of capsules and 300 µL of 10^-3^ TSB in PBS with or without 1 µg/mL ciprofloxacin (Fluka) were dispensed into wells. Additional 10^-3^ diluted TSB. with or without ciprofloxacin, was added at 4 h, 7 h, and 10 h of incubation at 37°C. In each subsequent addition, the ciprofloxacin concentration was adjusted to maintain a final concentration of 1 µg/mL in the media surrounding the capsules. For all time-lapse growth studies, the chambered coverglass was incubated at 37 °C in a microscope stage-top incubator (OKO Labs). Z-stacks of the capsules were imaged every 30-40 min using iCLSM (Leica).

**Figure 5:**
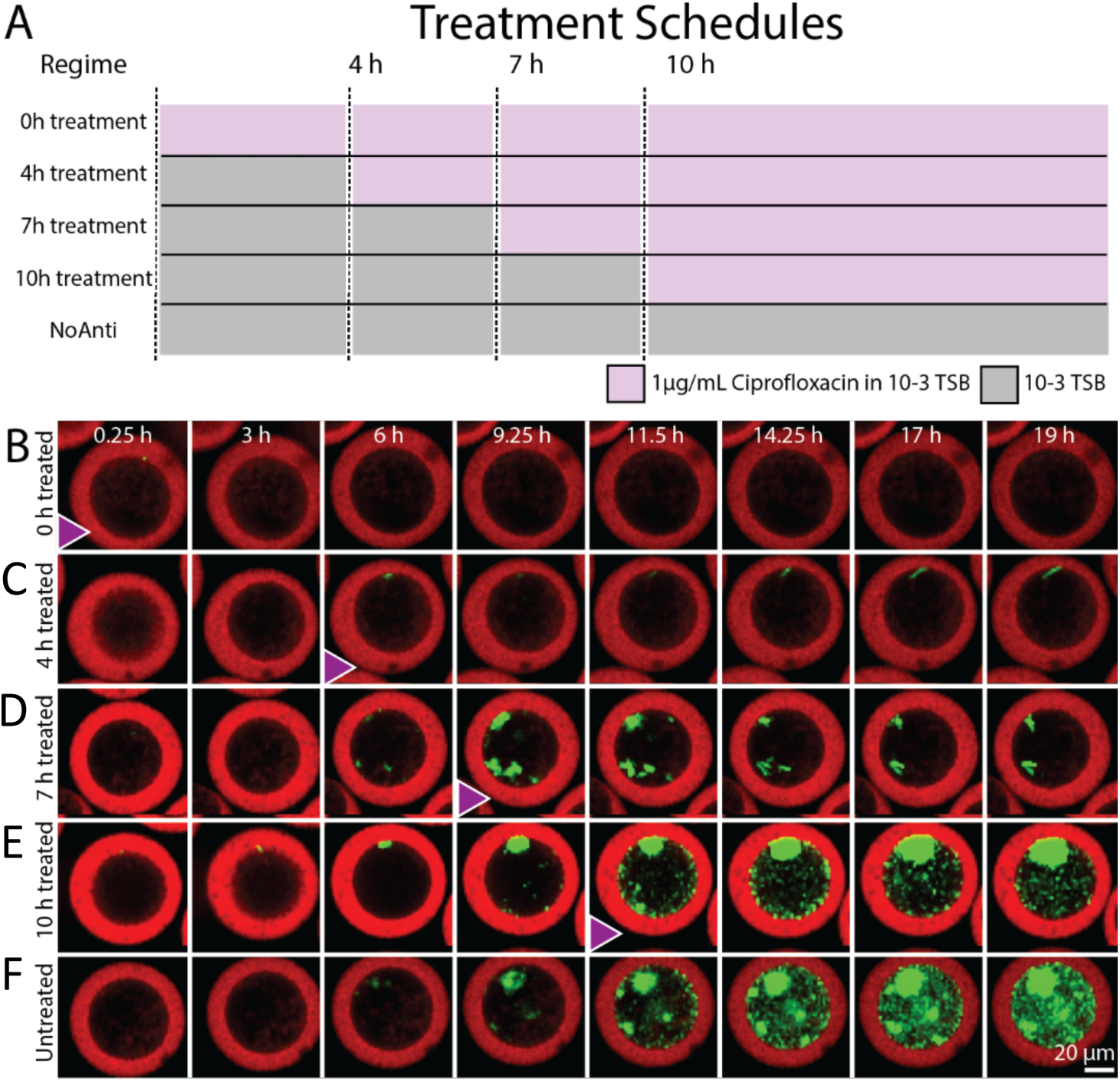
Antibiotic treatment of Bioflex encapsulated *P. aeruginosa*. (A) Capsules containing *P. aeruginosa* PAO1 were placed in the wells of a chambered coverglass with 10^-3^ TSB medium, with or without ciprofloxacin (1 µg/mL). (B-F) Ciprofloxacin was added to wells at 0, 4, 7, or 10 h after the start of incubation (addition point indicated by triangles), respectively. Bacterial growth within the capsules was imaged using iCLSM for each antibiotic treatment condition. The capsule shells were labelled with Cy5.5-maleimide (red), while *P. aeruginosa* expressed GFP (green).

**Figure 6:**
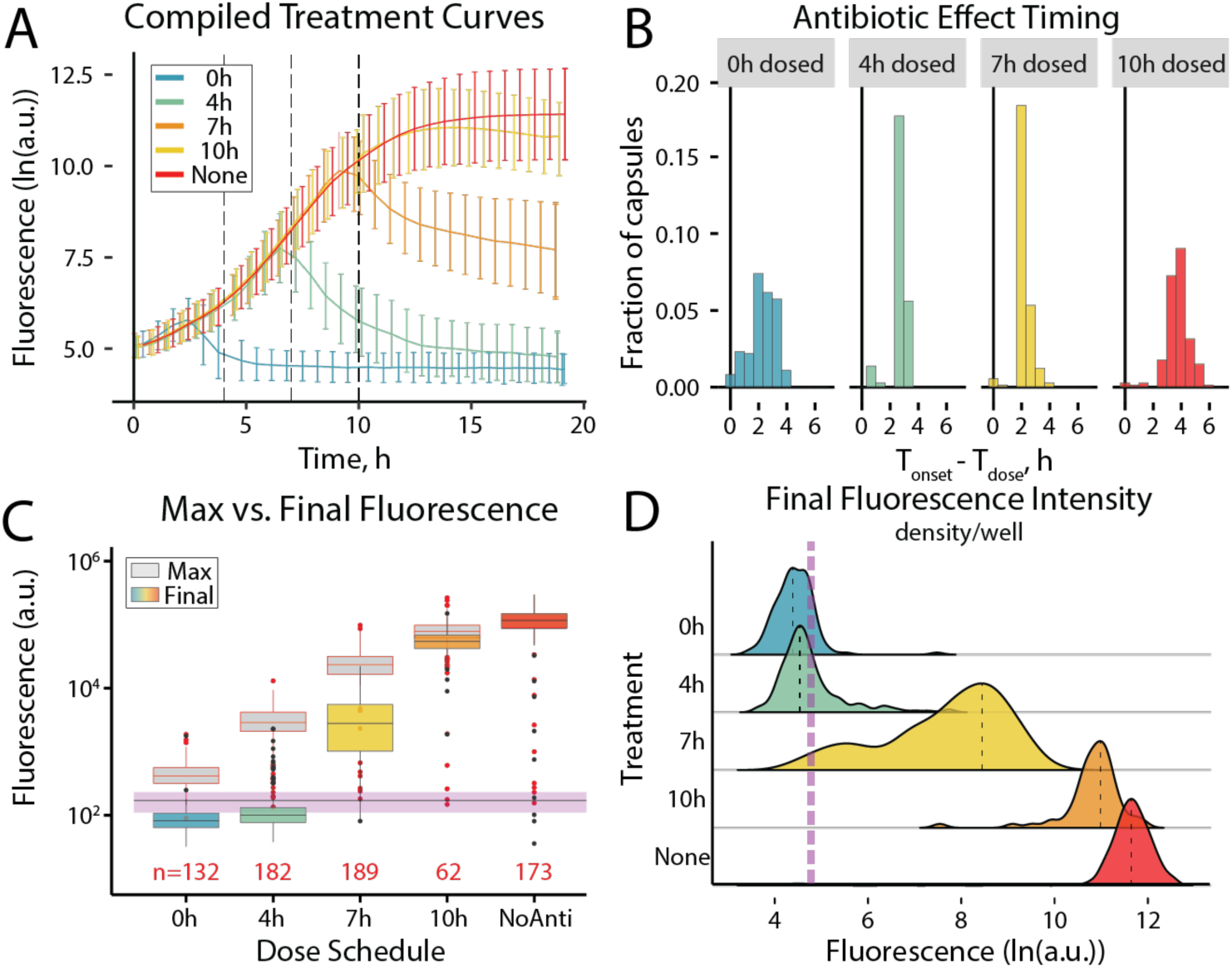
Heterogeneity in response of *P. aeruginosa* PAO1 cultured in Bioflex capsules starting with single cells. Treatment regimens correspond to Figure 6A. (A) The average growth and response to treatment, along with standard error, from bacteria cultured in capsules based on fluorescence intensities obtained from iCLSM z-stacks. (B) Histograms showing the time span between ciprofloxacin introduction and the onset of bacterial response, for each capsule in (A), indicating the timing of the treatment effect. (C) Box plot comparing initial fluorescence (purple line), maximum fluorescence (gray), and final fluorescence (colored) values for each treatment. The horizontal, dashed black line and purple bar represent the average fluorescence measurement and standard deviation at the initial time point. Boxes positioned below the purple line indicate that the final fluorescence was lower than the fluorescence output of a single cell, suggesting cell death. The n-value represents the number of bacteria-containing capsules analyzed. (D) Staggered density plots of total fluorescence intensity at the final time point for capsules exposed to each treatment. The vertical purple dashed line represents the average fluorescence of the compiled populations at the initial time point, while the black dashed lines within each density distribution indicate the mean fluorescence for that condition of that distribution. Data represent one biological sample, with multiple capsules analyzed per treatment regimen, as indicated in panel (C).

Bacterial growth within capsules (Figures 4-6, Supplementary Figures 3, 5, and 6) was quantified using fluorescence data from XYZ-T imaging stacks. Three biological replicates were analyzed to assess *P. aeruginosa* growth in TSB-containing capsules (Figure 4, Supplementary Figure 3), while a single biological sample was tested in the ciprofloxacin treatment assay (Figures 5, 6, Supplementary Figures 5,6). For each biological sample in either assay, 3-4 imaging fields of view were collected at 30-40 min intervals. Fluorescence imaging was performed using excitation and emission spectra corresponding to Cy5.5 and green fluorescent protein (GFP). The z-stack for each field of view at each time point was compressed into a single image using a summed projection of pixel intensities. Each image was intensity-thresholded to isolate usable capsule data. Usable data included cell-containing capsules where the entire capsule volume was captured in the z-stack and where fluorescent signal correlating to cell growth (GFP) from two different capsules did not overlap in the projected image. The ImageJ plugin TrackMate was used to quantify the fluorescence in each capsule at each time point. In the TSB growth assay, each field of view contained 5-54 cell-containing capsules that were analyzed for growth. In the ciprofloxacin treatment assay, each field of view contained 49-172 capsules. The variation in capsule number reflects differences in capsule concentration across fields of view. The resulting fluorescence data were imported into Rstudio for processing.

To subtract background fluorescence, the average z-projection summed fluorescence value from a sample of non-cell containing drops was calculated for each time point in each trial. This value was subtracted from the fluorescence intensity of cell-containing capsules. The background-subtracted, summed intensity values for each capsule were plotted as summed fluorescence vs. time on a semi-log plot. To determine the growth rates of TSB-only capsules (Figure 4, Supplementary Figure 3), fluorescence data were natural log-transformed for the period between 4.5 and 6 h, corresponding to log-linear growth. A linear regression was performed on per-capsule fluorescence intensity values against time to calculate the growth rate. The datasets for each trial were compiled and the mean growth rate value and standard deviation for capsule-based growth curves in TSB were calculated using a mixed effects model in RStudio, with biological replicate and the field of view as random effects.

To calculate the onset of ciprofloxacin’s effect, the ciprofloxacin delivery time was subtracted from the time point corresponding to the first observed fluorescence decrease in an individual capsule. A histogram of the effect of timing includes each individual capsule as an individual data point, with the averaged values reported in the text as simple means. All reported time values for the time-varied growth studies inherently includes a ± 40-min uncertainty, as response timing cannot be resolved below the 40-min imaging interval.

### 96 Well-Plate Growth Studies

Washed working cultures of *P. aeruginosa* PAO1, *P. aeruginosa* PAO1 pMF230, and *Escherichia coli* HB101 pMF230 ^31^ were prepared using the methods described in the "Preparation of Bacteria for Encapsulation" section. After a 30-min incubation, the optical density at 600 nm (OD_600_) of the washed cultures was measured using an Ultrospec 2100 Pro spectrophotometer (Amersham Biosciences). Cells were then diluted to an OD_600_ of 1.0 in PBS (pH 7.4), and 10 µL aliquots were used to inoculate the wells of sterile, black, clear bottom 96-well assay plates (Corning Incorporated) containing 190 µL media. The media conditions tested included PBS, TSB, 1:10 V/V 4- arm PEG-mal microspheres (described below) in PBS and in TSB, and 26 mg/mL dextran in PBS and in TSB, to evaluate the potential effects of capsule components on bacterial growth. All media conditions were amended with 150 µg/mL carbenicillin for strains carrying the pMF230 plasmid. For each media type and strain, three wells were inoculated as technical replicates. Positive growth control wells and negative sterile control wells (containing media without cells) were included in each assay. Microplates were lidded, sealed with Parafilm, and incubated at 37°C with constant shaking in a Cytation 5 Microplate Reader (BioTek Instruments). The OD_600_ and GFP fluorescence of each well were recorded every 15 minutes for 15 h. Three independent biological replicates were conducted for each condition.

The 4-arm PEG-mal microspheres, used to assess microbial consumption of capsule shell material, were synthesized using the same DTT nanoemulsion and 4-arm PEG-mal solution described in the "Microcapsule Components" section. The microspheres were generated as described in the "Microcapsule Generation" section, with one key modification: the microspheres lacked a core, and therefore, dextran solution was omitted from the capsule generation process. To achieve this, the core inlet of the microfluidic capsule generation device was plugged with a short section of sealed tubing. The flow rates used for microsphere generation were 150 µL/h 4-arm PEG-mal solution, 800 µL/h for the Span80/mineral oil solution, and 800 µL/h for the DTT nanoemulsion.

For dextran and 4-arm PEG-mal bulk growth studies, technical replicates were averaged and the mean and standard deviation of the three independent biological replicates were plotted using GraphPad Prism V10.3.1. Growth rates from cells within wells of microplates were calculated using the same approach as rates from cells within capsules. Briefly, background correction for optical density was performed by subtracting the average optical density reading from wells containing media only. For fluorescence background correction, the average fluorescence measurements of non-GFP producing *P. aeruginosa* control cells grown in wells was subtracted to correct for media autofluorescence. The background-subtracted values for each well were plotted against time on a semi-log plot. To determine growth rates, fluorescence data from the log-linear growth phase (between 2 and 3 h) were natural log-transformed, and a simple linear regression was performed in GraphPad Prism V10.3.1 to calculate the slope.

To assess the susceptibility of *P. aeruginosa* containing pMF230 to ciprofloxacin, broth microdilution assays were performed in TSB amended with 150 µg/mL carbenicillin using a Cytation 5 Microplate Reader (BioTek Instruments), following the Clinical and Laboratory Standards Institute (CLSI) standard operating procedure M07^32^ The minimum inhibitory concentration (MIC) was determined by applying a Gompertz model to the plot of log antibiotic concentration vs. the fractional area under the OD_600_ growth curve at each concentration in GraphPad Prism V9.5.1. The mean MIC and standard deviation of four independent biological replicates is reported.

### Data Analysis and Representation

All imaging data were processed using Fiji/ImageJ ^28^. Within ImageJ, the following plugins were used: the Hough circle transform ^29^ developed as part of the University of California, Berkeley Vision Sciences core grant (NIH P30EY003176) and TrackMate7 ^33^. Analysis and visualization of imaging data were performed in RStudio ^34^ using base R ^35^ along with the following packages: tidyverse^36^, ggplot2^37^, dplyr^38^, broom^39^, modelr^40^, scales^41^, lme4^42^, lmerTest^43^, and a ridgeline plot coded in base R (https://github.com/R-CoderDotCom/ridgeline).

For all boxplots, the elements are represented as follows: the median is shown as a bar; the interquartile range (*IQR*) spanning the 25^th^ to 75^th^ (Q1 to Q3) is shaded; the minimum and maximum values, calculated as 𝑄1 − 1.5 × 𝐼𝑄𝑅 and 𝑄3 + 1.5 × 𝐼𝑄𝑅, are represented by error bars; and outliers are shown as individual points. The median and quartile ranges are calculated directly for all boxplots, and do not represent data from mixed effects models.

## Results

### Generation of Bioflex Capsules: Hydrogel Selection and Crosslinking Strategy

The Bioflex method produces hydrogel-shelled capsules designed for single-cell bacterial isolation and cultivation. This process utilizes a non-planar microfluidic device to co-flow dextran and 4-arm PEG-maleimide solutions with mineral oil and a DTT nanoemulsion through microscale channels and channel junctions, generating hundreds of capsules per second (Figure 1A-B, Supplementary Figure 1). During capsule generation, the non-planar device creates an annular flow of dextran-rich core solution within the 4-arm PEG-mal shell precursor (Figure 1C). This flow intersects with mineral oil solution at a flow-focusing junction, forming aqueous drops in oil (water/water/oil emulsions, Figure 1A-B). Upon droplet breakup, the annular flow profile established by the 3-D device results in a radially heterogeneous droplet, where the 4-arm PEG-maleimide solution forms a shell-like layer surrounding the dextran-rich core.

The dextran–rich core and 4-arm PEG-mal shell precursor solutions form an aqueous two-phase system (ATPS), in which aqueous polymer solutions undergo phase separation when mixed at sufficiently high concentrations. The ATPS phase diagram delineates the mixture compositions that result in either a homogenous solution or a two-phase mixture. The boundary between these states is the binodal curve, where mixtures above the curve separate into two distinct phases, while those below the curve remain homogeneous (Figure 2A). Imaging of dextran and uncrosslinked 4-arm PEG-mal drops reveals the phase-equilibrium dynamics of these mixtures relative to the phase-separation curve. Above the curve, drops form a Janus particle structure, characterized by an eccentrically positioned dextran core surrounded by a 4-arm PEG-mal layer ^44^. Below the curve, drops form a homogenous solution containing both compounds (Figure 2B). Utilizing an ATPS for the core and shell precursor solutions during capsule generation ensures phase separation at the microfluidic junction prior to droplet breakup, leading to a well-defined boundary between the core and shell regions after shell crosslinking.

A channel carrying DTT nanoemulsion merges with the dextran in 4-arm PEG-mal drops downstream of the droplet breakup junction (Figure 1A-B). DTT rapidly partitions into the aqueous drops ^25,45^. The DTT thiol groups crosslink the 4-arm PEG-mal, solidifying the shell and preserving the core-shell geometry. Introducing DTT downstream of the drop generation junction prevents premature gelation at the junction, which could otherwise lead to device clogging ^21^.

After the addition of DTT, the stream of crosslinked capsules, mineral oil, and DTT nanoemulsion exits the device and is collected in a microcentrifuge tube for further processing (Figure 1A). The dimensions and geometries of the channels for the capsule-generating device are designed to minimize hydrodynamic resistance in the outlet channel while maximizing resistance in the core channel. Hydrodynamic resistance, calculated as the pressure drop for a given segment under capsule-producing flow rates with water as the flow medium, is lowest in the outlet channel and highest in the core channel (Supplementary Figure 1). This design prevents fluid backflow within the device, particularly into the aqueous channels, ensuring that DTT does not interact with 4-arm PEG-mal upstream, thereby avoiding device clogging.

#### Control of Capsule Shell Thickness using Flow Rate

The Bioflex method enables precise control over capsule shell thicknesses by adjusting flow rates without altering the capsule formulation. By decreasing the 4-arm PEG-mal volumetric flow rate from 80% to 30% of the total aqueous flow (*Q_aq_*), while maintaining a constant overall aqueous flow rate (Supplementary Table 1), intact capsules were generated with shell thicknesses ranging from 28.1 ± 2.0% to 71.5 ± 0.9% of the total capsule radius (Figure 3A, 3B). Core-shell capsules with well-defined core and shell regions were successfully produced at 4-arm PEG-mal flow rates as low as 10% of the total aqueous flow.

However, when the 4-arm PEG-mal flow rate fell 30%, capsules exhibited poor structural integrity and consistency, often failing to form fully enclosed, spherical capsules (Figure 3A, 3C, points 5-6). The specific flowrates used for each particle are listed in Supplementary Table 1.

#### Control of Capsule Size using Flow Rate

As with shell thickness, the capsule radii can be tuned by modifying flow rates during capsule generation (Figure 3D, 3E).

Decreasing the combined aqueous flow from 17% to 3% of the overall flow resulted in capsule radii ranging from 45.9 ± 1.3 μm to 29.5 ± 2.3 μm. The specific flow rates used to generate capsules of each size are detailed in Supplementary Table 2.

#### Capsules are Generated Below Phase-Separating Concentrations of 4-arm PEG-mal and Dextran

To assess the capability of the Bioflex method to generate capsules at 4-arm PEG-mal and dextran concentrations below the ATPS binodal curve, the composition of Bioflex generated core-shell capsules was compared to the phase separation diagram for 4-arm PEG-mal and dextran (Figure 3C). Adjusting the ratio of 4- arm PEG-mal solution to dextran solution during capsule generation resulted in a range of total capsule concentrations that spanned both the phase-separating and non-phase separating regions of the diagram (Figure 3C). The Bioflex method successfully induced phase separation and shell gelation under all tested flow conditions, including cases where the 4-arm PEG-mal flow rate was less than 50% of the total flow rate. This corresponded to in-capsule concentrations of 4-arm PEG-mal and dextran that fell within the non-phase-separating region of the phase separation diagram (Figure 3 A, 3C), below the binodal curve.

#### Growth of Bacteria in Capsules Starting from Single Cell Inocula

We encapsulated single *P. aeruginosa* (pMF230) cells in Bioflex capsules to assess their potential for measuring bacterial growth kinetics and to determine whether the capsule components affect cell viability. Capsules containing single cells were suspended in TSB medium within the wells of a chambered coverglass, incubated for 12 h at 37 °C, and imaged at 30-min intervals by iCLSM. The images show an increase in cell-associated fluorescence inside the capsules over time (Figure 4A). The total background-subtracted fluorescence per capsule was quantified from the imaging data and plotted over time (Figure 4B). The growth rate within each individual capsule was calculated from the exponential portion of the growth curve (Figure 4C). The results indicate that the average growth rate of capsule-cultivated *P. aeruginosa* is 1.08 ± 0.031 h^-1^. We quantified the growth rates starting from 151 single cells, revealing variations in doubling time among individual cells.

To determine whether incubation in Bioflex capsules influences cellular growth rates, we also quantified *P. aeruginosa* (pMF230) growth in TSB using a microplate reader. Growth was monitored via optical density (OD_600_) and GFP fluorescence over time. Using this conventional approach, the growth rate was determined to be 1.25 ± 0.041 h^-1^, based on OD_600_, and 1.19 ± 0.039 h^-1^ based on fluorescence (Supplementary Figure 4). The doubling times measured by fluorescence and OD_600_ were not significantly different (*p* = 0.125, unpaired t-test). These results suggest that while fluorescence may be influenced by GFP expression, maturation time, or plasmid loss, it remains a reliable metric for estimating growth under these conditions. Furthermore, the growth rate measured in capsules was not significantly different from that obtained using the bulk method. Together, these findings demonstrate that changes in bacterial fluorescence within Bioflex capsules can be used to monitor growth parameters from single *P. aeruginosa* cells. A key advantage of the capsule-based assay over traditional well-plate methods is that it provides insights into the range of growth rates among individual cells, allowing for quantification of population heterogeneity (Figure 4C). Interestingly, iCLSM of *P. aeruginosa* in capsules revealed that *P. aeruginosa* microcolonies preferentially form along the inner interface of the capsule shells, suggesting the potential for biofilm formation on capsule walls. This observation highlights the utility of capsule-based growth assays not only for measuring growth rates, but also for studying of colony formation and cell morphology.

#### *P. aeruginosa* Does Not Utilize Capsule Components for Growth

4-arm PEG-maleimide and dextran are carbon-containing polymers and could potentially serve as carbon sources for bacteria during capsule-based growth assays, which could limit the assay’s ability to test specific nutrient conditions. To assess whether these capsule components can serve as a carbon source, *P. aeruginosa* (pMF230) cells were incubated with 10% crosslinked 4-arm PEG-mal or 26 mg/ml dextran to determine whether these compounds are degraded by bacteria, and could thus influence bacterial growth in capsules (Supplementary Figure 4). Growth, monitored as O.D._600_, did not occur in PBS over 15 h, even with the inclusion of 4-arm PEG-mal or dextran (Supplementary Figure 4). The addition of these capsule components to TSB did not increase the growth of PAO1 pMF230 cells over 15 h. *Escherichia coli* HB101 (pMF230) cells were also monitored under these conditions and were not affected by the presence of either 4-arm PEG-mal or dextran. These results demonstrate that capsule components do not significantly influence bacterial growth of these two microorganisms within the timeframe of these experiments.

#### Encapsulated *P. aeruginosa* Responds to Changes in the Extracapsular Environment

The Bioflex capsules are designed to allow the diffusion of nutrients and other chemicals between the external environment and the capsule interior. Therefore, modifying the external environment should enable the observation of cellular responses within the capsules. To determine whether Bioflex capsules permit the diffusion of chemicals such as antibiotics, we exposed encapsulated *P. aeruginosa* to ciprofloxacin, a DNA gyrase-inhibiting antibiotic, by altering the external environment. Single *P. aeruginosa* cells were encapsulated in Bioflex capsules and cultured in a chambered well-slide with 10^-3^ diluted TSB media for 20 h. The diluted TSB was used to limit culture density in the capsules, restrict the proliferation of escaped (extracapsular) cells, and maintain high-quality microscopy over an extended time frame. Ciprofloxacin (1 µg/mL in 10^-3^ TSB) was introduced at 0, 4, 7, or 10 h of incubation on the slide (Figure 5A).

This inhibitory concentration was selected based broth microdilution MIC assays which determined that the MIC of PAO1 pMF230 in TSB was 0.26 ± 0.05 µg/mL. Capsules were imaged using iCLSM at 40 min intervals over a 19.5 h incubation period (Figure 5B-F, Supplementary Figure 5). Time-lapse images (Figure 5B-F and Supplementary Figure 5) revealed an increase in fluorescent signal per capsule for all samples before antibiotic addition, except for those treated with ciprofloxacin at 0 h. As shown in Figure 5D-F, cell aggregates appeared as dense microcolonies associated with the capsule walls, while planktonic cells appeared as disperse, punctate fluorescent foci throughout the capsules. The time-lapse image series indicated differences in morphology and growth patterns depending on the growth phase and the number of cells contained in each capsule at the time of antibiotic addition (Figure 5B-F, Supplementary Figure 5).

In the continuous antibiotic treatment condition (0 h treated well), no growth was observed, indicating that ciprofloxacin diffused into the capsules. When ciprofloxacin was added at 7 h of growth, fluorescence initially increased before decreasing (Figure 6A, Supplementary Figure 6C). When antibiotics were introduced after 10 h of growth, fluorescence did not noticeably decrease, but additional bacterial growth appeared inhibited (Figure 6A, Supplementary Figure 6D). Encapsulated cells left untreated for 19.5 h showed no reduction in fluorescence (Figure 6A, Supplementary Figure 6E). The results demonstrate that Bioflex capsules are permeable to ciprofloxacin and suggest that the growth phase and microcolony formation influence the response of *P. aeruginosa* to antibiotic treatment.

#### Cell Heterogeneity in Response to Antibiotic Treatment

The Bioflex capsules allow quantification of heterogeneity in the response of single cells or microcolonies originating from single cells to antibiotic treatments. (Figure 6A). Plots of increase/decrease in fluorescence per capsule are shown in Supplementary Figure 6. Bacterial fluorescence measurements from each sample demonstrate that treating encapsulated cells at different time points resulted in distinct effects (Figure 6, Supplementary Figure 6). Fluorescence values reported in Figure 6A represent the average and standard error of the per-capsule total intensity for each cell or microcolony, and are normalized to the total fluorescence value at *t* = 0 h. Continuous treatment with ciprofloxacin at 4X the MIC produced a significantly different growth outcome compared to untreated capsules. In untreated capsules, fluorescence increased log-linearly from 2 to 12 h before reaching stationary phase. By 19.5 h, the average fluorescence per capsule was 1.08 x 10^5^ ± 2.47 x 10^3^ arbitrary units (a.u.). In contrast, fluorescence from continuously treated bacteria initially increased to 97.5 ± 18.4 a.u. at 2.5 h, then declined to a final value of -65.5 ± 14.3 a.u. (Figure 6A). All reported increases and decreases in total fluorescence intensity were statistically significant at a 95% confidence level (α = 0.05) (Supplementary Table 4).

The onset of antibiotic-effect was determined by identifying the first negative slope in fluorescence curves between adjacent time points ((Figure 6A, 6B. Supplementary Figure 6). No negative slopes were observed in untreated capsules. An antibiotic effect was detected at 2 - 2.5 h post-ciprofloxacin introduction for the 0, 4, and 7 h treated samples, and at 4 h for the 10 h treated samples (Figure 6B).

Bacteria in capsules treated with ciprofloxacin at 4, 7, and 10 h exhibited heterogeneous responses, with outcomes ranging between those of untreated and continuously treated cells. Figure 6C displays the variation in treatment responses across 738 capsules containing cells or microcolonies under five treatment conditions. Continuously treated samples (*n* = 132) initially showed increased fluorescence before declining below the initial inoculum level. For the 4 h treated capsules, fluorescence initially increased until 6.5 h (1.62 x 10^3^ ± 46.8 a.u.), then declined by four orders of magnitude to -7.36 ± 1.8 a.u (*n* = 182), with some outliers. Bacteria treated following 7 h of growth exhibited the greatest range in responses. Fluorescence peaked at 9 h (1.78x10^4^ ± 1.29x10^2^ a.u.) before decreasing by one order of magnitude to a 2.77x10^3^ ± 90 a.u. (*n* = 189). The wide range of responses (yellow box, Fig 6C) indicates the most heterogeneity in responses for these encapsulated microcolonies. The 10 h treated sample showed the least response to treatment but included outliers (Figure 6C) (*n* = 62). Fluorescence for this sample group peaked at 14 h (7.33 x 10^4^ ± 1.02 x 10^2^ a.u.), before decreasing by less than an order of magnitude (5.51x10^4^ ± 1.66 x 10^2^ a.u).

The staggered density plot shows the range of responses of encapsulated *P. aeruginosa* to 1 µg/mL ciprofloxacin treatment at the final timepoint (Fig 6D). Encapsulated bacteria treated at 4 h and 7 h showed the greatest diversity in response to treatment. The 0 h treated and 10 h treated samples showed less diversity, but contained distinct outliers, suggesting higher tolerance to antibiotic treatment. Overall, these results demonstrate that *P. aeruginosa* growth within Bioflex capsules can be quantitatively analyzed and that the capsules are permeable to antibiotics. By tracking the responses of single cells and microcolonies derived from single cells in Bioflex capsules, the heterogeneity in response to antibiotics can be quantified using time-lapse imaging.

## Discussion

### Permeable Hydrogel Capsules for Optimizing Studies of Single Cell and Clonal Bacterial Lineages

This study aimed to develop permeable microfluidics growth chambers for single-cell studies and to establish a foundation for complex, multi-step microbial assays. The Bioflex capsule generation method produces permeable, biocompatible, and structurally tunable hydrogel-shelled microcapsules isolating and cultivating single microbial cells. This method optimizes capsule production for microbial growth assays by integrating three key technologies: (1) an aqueous two-phase system (ATPS), (2) a non-planar microfluidic device, and (3) nanoemulsion-delivered crosslinking agent.

In non-ATPS based approaches, core-shell mixing makes capsule formation highly sensitive to crosslinking timing ^25,46–48^. Additionally, because the core and shell solutions of non-ATPS systems start mixing upon contact, the boundary between the core-shell regions is less distinct than in ATPS-based capsules ^25,46^. The use of ATPS in the Bioflex method reduces the need for rapid crosslinking and enhances capsule geometry through phase separation.

ATPS-based capsule formation in planar microfluidic devices requires time for phase separation and reorientation into the desired core-shell geometry after drop generation ^20,21,49^. Capsule production is only possible using ATPS concentrations above the phase separation line, and delaying crosslinker introduction or slowing the kinetics of crosslinking is required to allow for an optimal amount of phase separation^20,21,49^. Thus, crosslinking must be carefully timed. If crosslinking occurs too quickly, the phases cannot fully reorient into a concentric core-shell structure; if crosslinking occurs too slowly, phase separation can progress into an undesirable eccentric Janus particle configuration^44^.

The Bioflex method addresses these limitations by using a non-planar device geometry and channel configuration, similar to those used for non-ATPS capsule production ^25,26^, to control phase separation and orientation before droplet breakup. This results in precise, flow-controlled templating of the core-shell structure. Additionally, nanoemulsion-delivered DTT crosslinking in conjunction with the 3-D device rapidly freezes the capsule geometry after drop formation, crosslinking the templated geometry. The method also enables capsule generation with 4-arm PEG-maleimide and dextran concentrations below the binodal curve, as localized phase separation at the droplet interfaces supports core-shell formation before rapid crosslinking.

### The Bioflex Capsule Generation Method Tolerates Widely Varied Operating Dynamics

Shell thickness and capsule radius results indicate that the Bioflex method is compatible with a broad range of flow rates and dextran or 4-arm PEG-mal concentrations. This flexibility, combined with a clog-resistant device design, simplifies operation, and enables varied particle geometries without requiring precursor reformulation. By adjusting shell thickness and capsule size, the method could be leveraged to investigate spatial dependencies in cell-to-cell signaling ^12^ and polymicrobial interaction ^50^, as well as fine-tune multi-capsule neighborhood dynamics. The method also allows for customization of core and shell compositions to meet experimental needs without disrupting capsule formation. However, some capsules exhibited eccentric core positioning, where shell thickness varied asymmetrically. A potential solution is density-matching the core and shell solutions to minimize buoyancy differences^20^, or modifying the channel junction to improve annular flow symmetry before droplet breakup.

### Bioflex Capsules Support Single Cell Growth Assays

PEG-4-mal has been widely used in cell culture, confirming its biocompatibility ^21,25,26,51^. Here, we demonstrated that *P. aeruginosa* growth rates in PEG-4-mal capsules were comparable to traditional 96 well-plate assays. By culturing *P. aeruginosa* and *E. coli* in well plates in the presence and absence of capsule components, we confirmed that the capsule components alone do not impact bacterial growth (Supplementary Figure 4). For long-term in-capsule cultivation, mutant screenings, or starvation studies, it may be necessary to reassess bacterial interactions with capsule components over extended exposure.

Another consideration using hydrogel capsules for microcultures is cell containment. Previous studies utilizing microspheres instead of capsules require processes such as particle coating or surface sterilization to prevent contamination of the bulk media by escaping bacteria ^52,53^. In this work, intact capsule shells effectively contained cells, as no leakage was observed during extended cultivation times.

However, robust bacterial growth within capsules did sometimes lead to rupture and cell release, a phenomenon also reported in prior studies ^21^. Prevention of capsule overgrowth and rupture can be achieved by reducing the nutrient concentrations, as demonstrated in our study using 10^-3^ diluted TSB, where no rupture was observed.

Additionally, densely packed capsules can limit the ability to perform quantitative fluorescence analysis using light microscopy. Optimizing the uniformity of capsule shell thickness or tuning the capsule shell composition may improve capsule resilience to rupture. Other research indicates that 4-arm PEG-maleimide capsules can swell ^21^ to accommodate an increasing volume of cells as cells grow. We also observed increased capsule size for capsules full of cells in some instances. Tuning the composition of the cross-linkable hydrogel used in hydrogel capsule production to improve mechanical properties, including capsule elasticity, durability, and structural symmetry, may further enhance the utility of capsules for microbial cultivation.

### Bioflex Capsules Enable Multistep, Single-Cell Assays, and Rich Data Output

The permeability of Bioflex capsules allows integration of multistep protocols for single-cell compartmentalization, and can facilitate studies on medically and environmentally relevant topics such as changes in the chemical environment ^54^, effects of antimicrobial dosing ^55,56^, and responses to nutrient limitation ^57^. Here, we demonstrated that Bioflex capsules enable long-term tracking of individual cells across multiple media exchanges. Additionally, water-soluble compounds such as ciprofloxacin can diffuse into capsules and affect the physiology of *P. aeruginosa,* demonstrating the adaptability of the capsule method for multistep assays.

Our findings revealed key factors in bacterial response to antibiotics, including microcolony morphology, heterogeneity of bacterial microcolonies in individual capsules, the heterogeneity of antibiotic treatment outcomes, and timing between antimicrobial exposure and observable effects. Other studies have shown that 4-arm PEG-mal microspheres and capsules are amenable to introducing dyes and other compounds useful in microbial analysis ^20^ and fluorescence-activated cell sorting (FACS) ^21^, further demonstrating the use of permeable, hydrogel-shelled capsules for multistep assays and downstream analysis. Future improvements could focus on optimizing media replacement strategies while maintaining capsule integrity and tracking.

### Bioflex Capsules as a Model for Microscale Microcosms

Bacteria cultured in Bioflex capsules exhibited phenotypic and morphologic complexity, with dense aggregates at the capsule interface coexisting with planktonic cells, all arising from a single-cell inoculum. Previously, our group studied the same *P. aeruginosa* PAO1 pMF230 strain in water-in-oil drops ^14,15^ and found that under identical nutrient conditions, bacterial growth in drops resulted in dispersed, planktonic growth.

This suggests that complex, aggregate-like colony morphology may be influenced by surface interactions or varying nutrient dynamics.

A major advantage of the Bioflex method is its potential for high-throughput single-cell assays. Capsules can be generated at kilohertz rates, and hundreds can be imaged simultaneously, enabling large-scale microbial growth studies. Unlike water-in-oil assays ^14,15^, which require separate sample preparation for each condition, a single batch of Bioflex capsules can be divided post-generation to test multiple conditions.

Large single-cell resolved sample sizes are essential for understanding microbial heterogeneity, as rare phenotypes are frequently responsible for medically-impactful community traits, for example, the role of persister cells in infection recalcitrance ^58^. Thus, single-cell capsule cultivation provides a powerful approach for identifying phenotypically diverse subpopulations at high sample sizes. The ability to track single cells over time under transient chemical conditions expands the scope of single-cell studies to systems with complex, time-dependent parameters.

## Conclusion

The results presented here demonstrate the utility of capsule-based cultivation for studying time-dependent antibiotic effects, providing high-resolution data on bacterial growth, treatment response, and effect timing. The Bioflex capsule platform offers a robust, scalable method for single-cell microbial cultivation, supporting long-term, time-resolved studies in diverse experimental conditions. By integrating high-throughput generation, controlled permeability, and multistep assay compatibility, Bioflex capsules provide a powerful tool for investigating microbial heterogeneity, antibiotic resistance, and cellular dynamics in response to environmental changes.

## SUPPLEMENTARY MATERIALS

### Supplementary Tables

**Supplementary Table 1:**
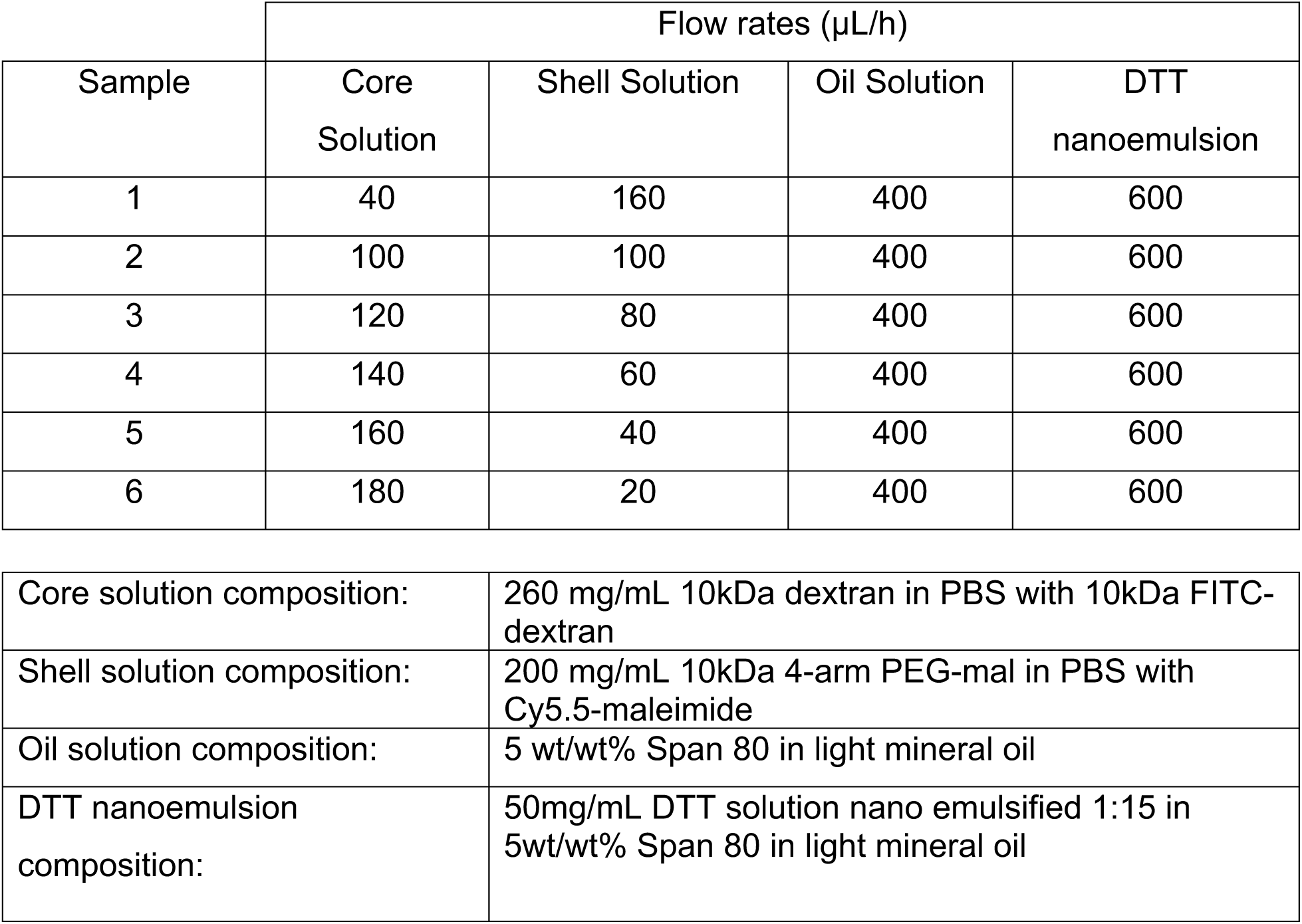
Capsule Shell Thickness Assay Flow Rates.

| Sample | Flow rates ($\mu\text{L/h}$) | | | |
| --- | --- | --- | --- | --- |
|  | Core Solution | Shell Solution | Oil Solution | DTT nanoemulsion |
| 1 | 40 | 160 | 400 | 600 |
| 2 | 100 | 100 | 400 | 600 |
| 3 | 120 | 80 | 400 | 600 |
| 4 | 140 | 60 | 400 | 600 |
| 5 | 160 | 40 | 400 | 600 |
| 6 | 180 | 20 | 400 | 600 |

|  |  |
| --- | --- |
| Core solution composition: | 260 mg/mL 10kDa dextran in PBS with 10kDa FITC-dextran |
| Shell solution composition: | 200 mg/mL 10kDa 4-arm PEG-mal in PBS with Cy5.5-maleimide |
| Oil solution composition: | 5 wt/wt% Span 80 in light mineral oil |
| DTT nanoemulsion composition: | 50mg/mL DTT solution nano emulsified 1:15 in 5wt/wt% Span 80 in light mineral oil |

**Supplementary Table 2:**
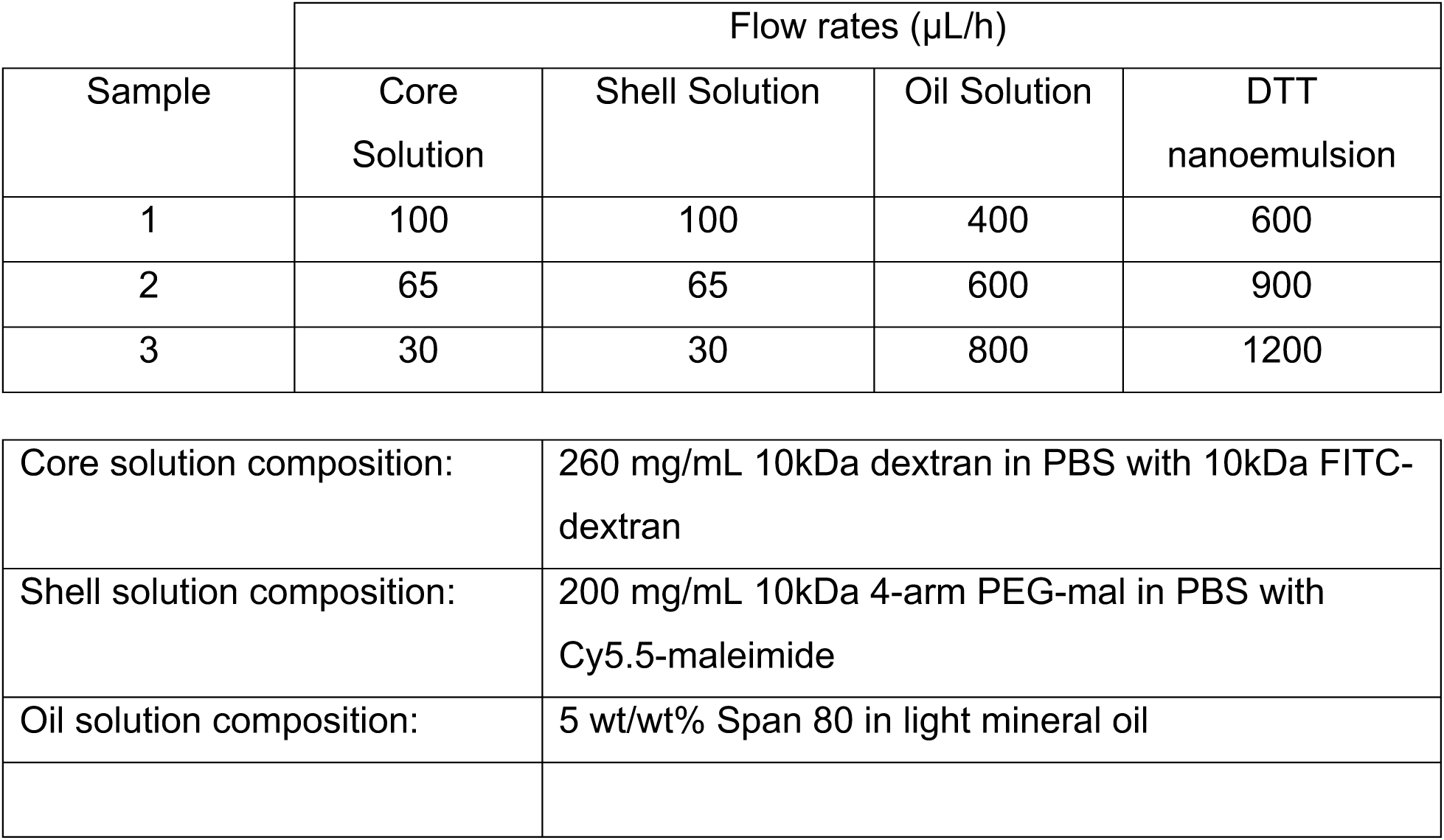
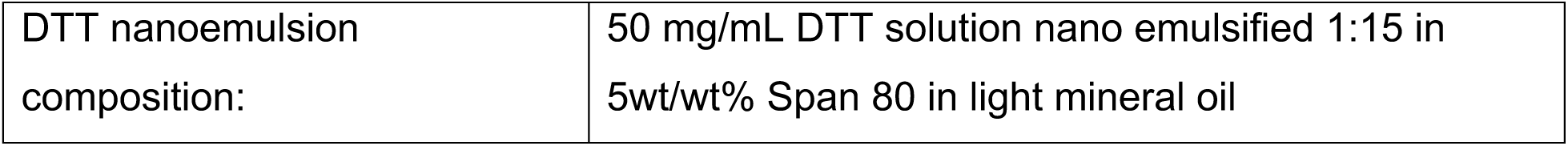
Capsule Radii Assay Flow Rates.

| Sample | Flow rates ($\mu\text{L/h}$) | | | |
| --- | --- | --- | --- | --- |
|  | Core Solution | Shell Solution | Oil Solution | DTT nanoemulsion |
| 1 | 100 | 100 | 400 | 600 |
| 2 | 65 | 65 | 600 | 900 |
| 3 | 30 | 30 | 800 | 1200 |

|  |  |
| --- | --- |
| Core solution composition: | 260 mg/mL 10kDa dextran in PBS with 10kDa FITC-dextran |
| Shell solution composition: | 200 mg/mL 10kDa 4-arm PEG-mal in PBS with Cy5.5-maleimide |
| Oil solution composition: | 5 wt/wt% Span 80 in light mineral oil |
| DTT nanoemulsion<br>composition: | 50 mg/mL DTT solution nano emulsified 1:15 in<br>5wt/wt% Span 80 in light mineral oil |

**Supplementary Table 3:**
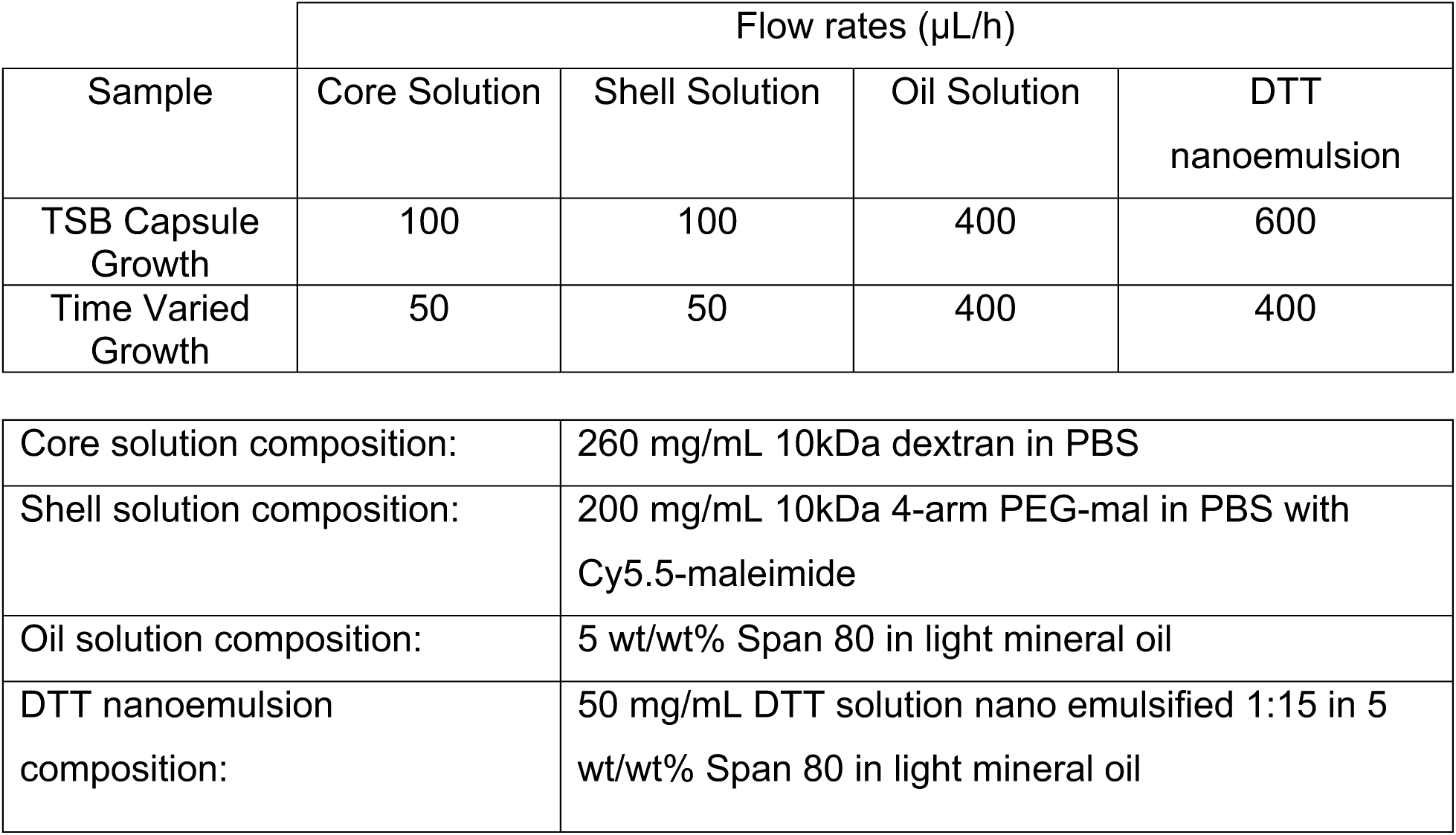
Biological Experiment Flow Rates.

| Sample | Flow rates ($\mu\text{L/h}$) | | | |
| --- | --- | --- | --- | --- |
|  | Core Solution | Shell Solution | Oil Solution | DTT nanoemulsion |
| TSB Capsule Growth | 100 | 100 | 400 | 600 |
| Time Varied Growth | 50 | 50 | 400 | 400 |

|  |  |
| --- | --- |
| Core solution composition: | 260 mg/mL 10kDa dextran in PBS |
| Shell solution composition: | 200 mg/mL 10kDa 4-arm PEG-mal in PBS with Cy5.5-maleimide |
| Oil solution composition: | 5 wt/wt% Span 80 in light mineral oil |
| DTT nanoemulsion composition: | 50 mg/mL DTT solution nano emulsified 1:15 in 5 wt/wt% Span 80 in light mineral oil |

**Supplementary Table 4:** Comparison of Means for Antibiotic Assay.

**Two Sample Comparison of Means (2 Tailed *t*-Test) *p* Values**
| Treatment | *T0* vs. *Tmax* | *T0* vs. *Tf* | *Tmax* vs. *Tf* | Sample Size |
| --- | --- | --- | --- | --- |
| 0 h | <2.2e-16 | 3.1E-08 | <2.2e-16 | 132 |
| 4 h | <2.2e-16 | 0.011 | <2.2e-16 | 182 |
| 7 h | <2.2e-16 | <2.2e-16 | <2.2e-16 | 189 |
| 10 h | <2.2e-16 | <2.2e-16 | 2.1E-05 | 62 |
| untreated | NA | <2.2e-16 | NA | 173 |

### Supplementary Figures

**Supplementary Figure 1:**
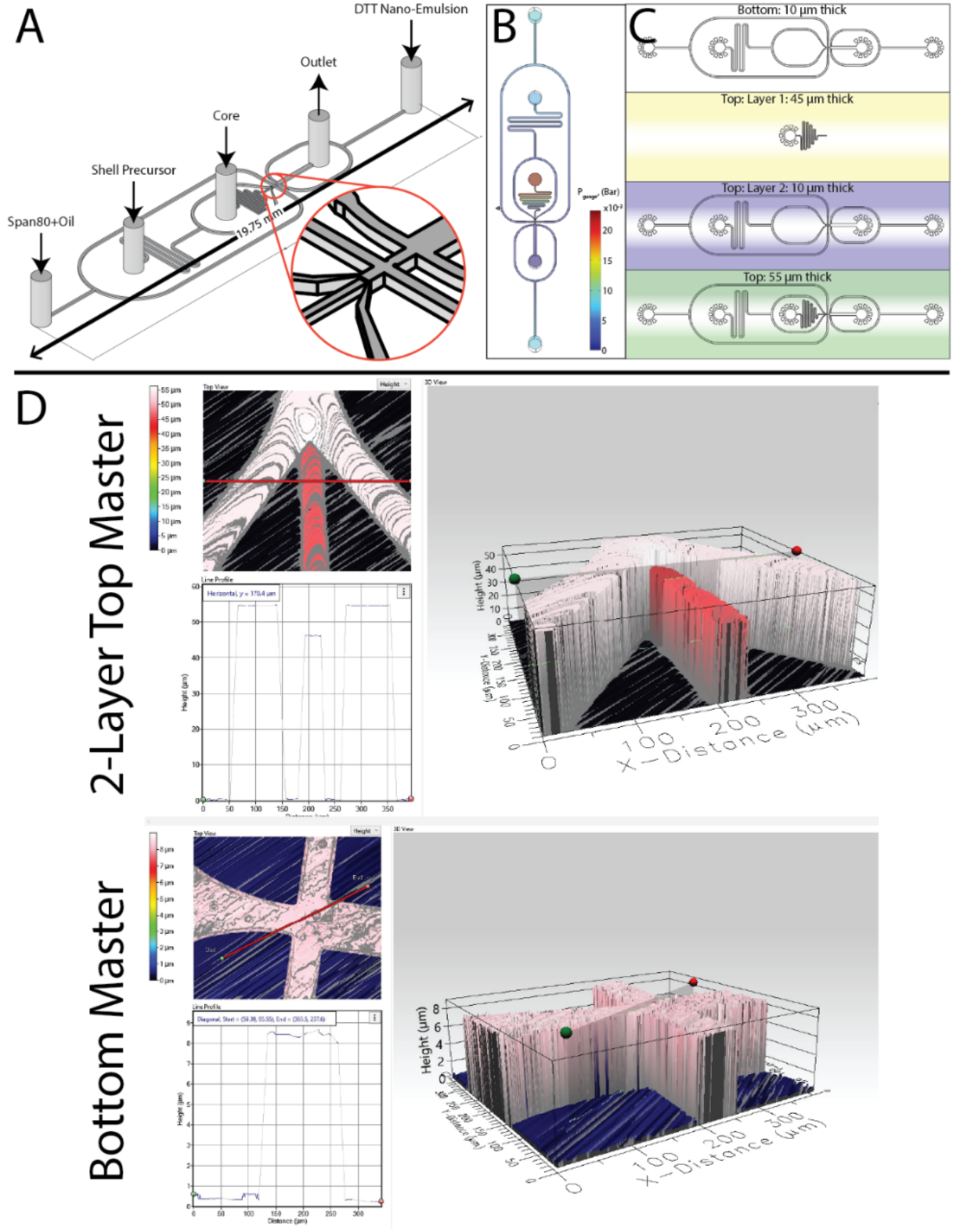
Bioflex capsule generation device design and fabrication notes. (A) The capsule generation device is a flow-focusing drop generation device with non-planar geometry at the junction of the core and sheath precursor channels. (B) A snapshot of pressure drop across the device for STP water at capsule-generation flow rates. Color gradient indicates pressure. (C) Non-planar geometry is achieved using multi-layered PDMS masters. The masters are made on silica wafers using 2 layers of photoresist on the top master and a single layer on the bottom master. (D) Profilometry was used to ensure master features are high fidelity and the desired height.

**Supplementary Figure 2:**
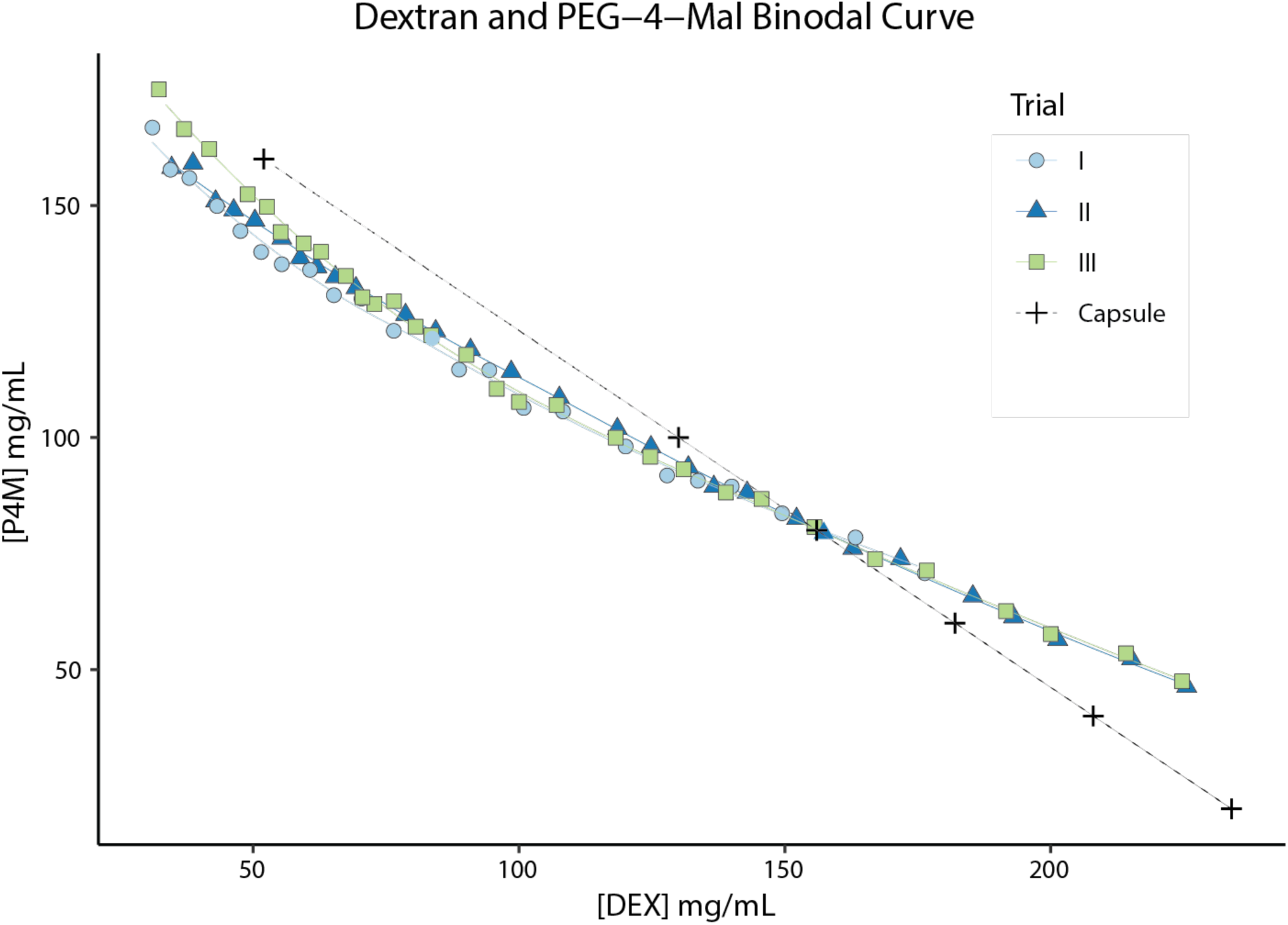
Individual cloud point titration trials used to determine the binodal curve of 4-arm PEG-mal / dextran ATPS used in the Bioflex capsules. Dashed line with + symbols represents the concentrations tested during the shell thickness variation experiment.

**Supplementary Figure 3:**
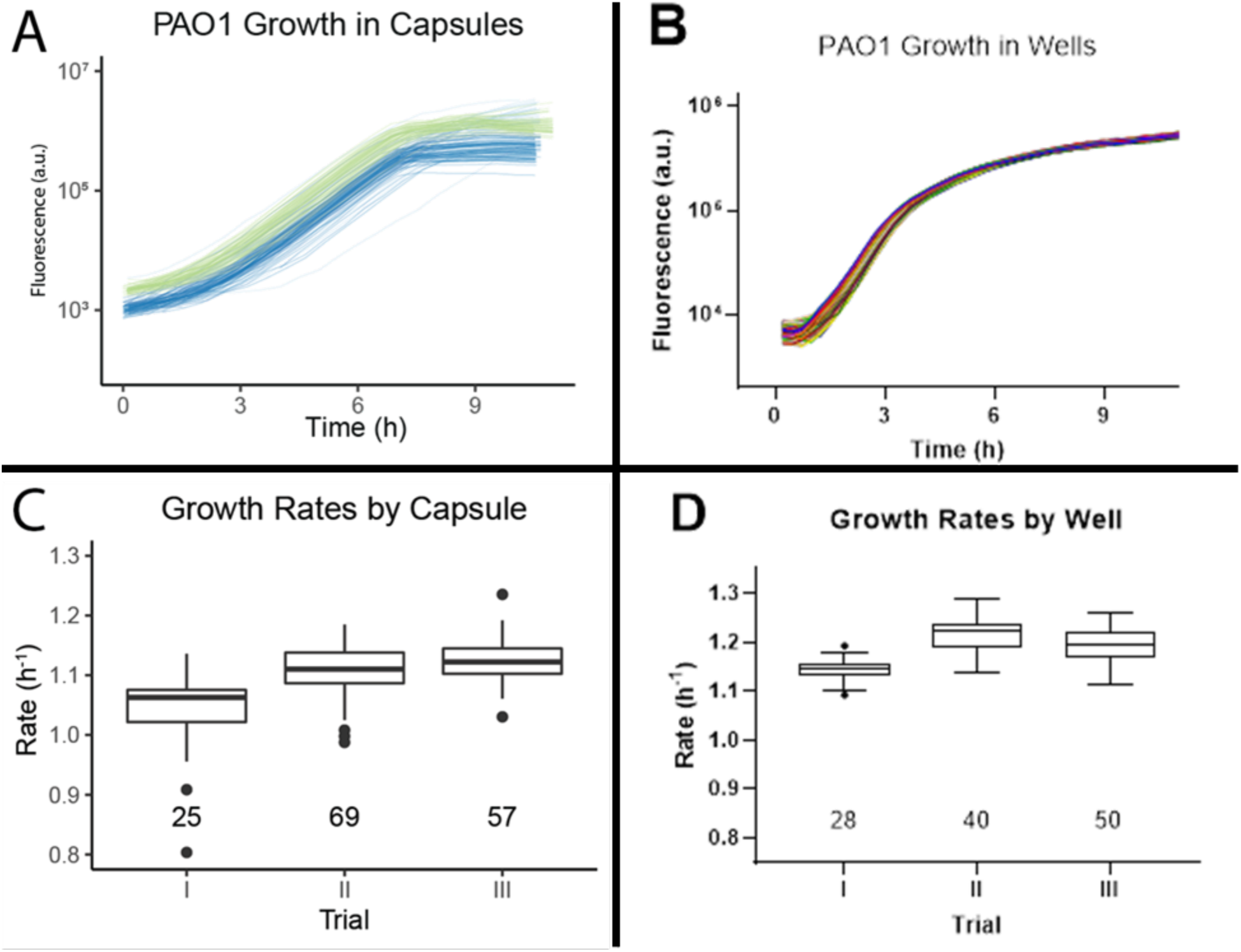
Summary of growth in capsules vs. bulk experiments. (A) Growth curves for bacteria in individual capsules or (B) individual wells of a 96-well plate. (C) Growth rates based on fluorescence increase for capsule trials and (D) for multiwell plate reader trials. Three independent biological experiments were performed for each dataset; n values per trial are listed directly above the trial number in panels C and D.

**Supplementary Figure 4:**
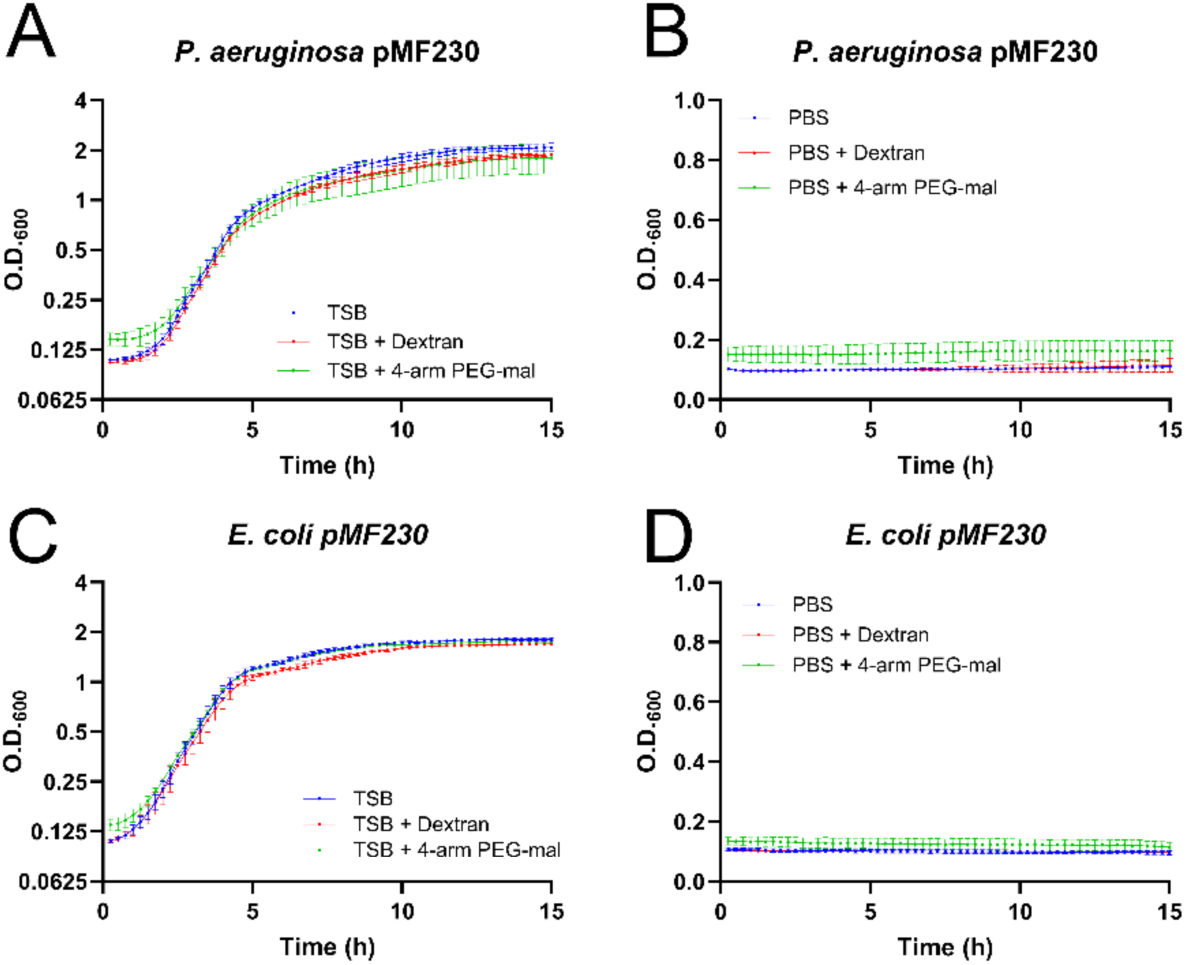
Optical density was monitored over 15 h in a microplate reader to determine the effect of capsule components on bacteria harboring the pMF230 plasmid. (A) *P. aeruginosa* PAO1 (pMF230) cells and (C) *E. coli* HB101 (pMF230) cells did not exhibit a significant increase or decrease in growth when 4-arm PEG-mal or Dextran were added to the TSB growth medium. Additionally, no effect was observed when 4-arm PEG-mal or Dextran were added to (B) *P. aeruginosa* PAO1 (pMF230) cells in PBS medium lacking nutrients or to (D) *E. coli* HB101 (pMF230) cells in PBS medium. The average and standard deviation of three independent biological replicates (calculated from three technical replicates each) are plotted, demonstrating that the capsule components do not significantly affect the proliferation of these two strains.

**Supplementary Figure 5:**
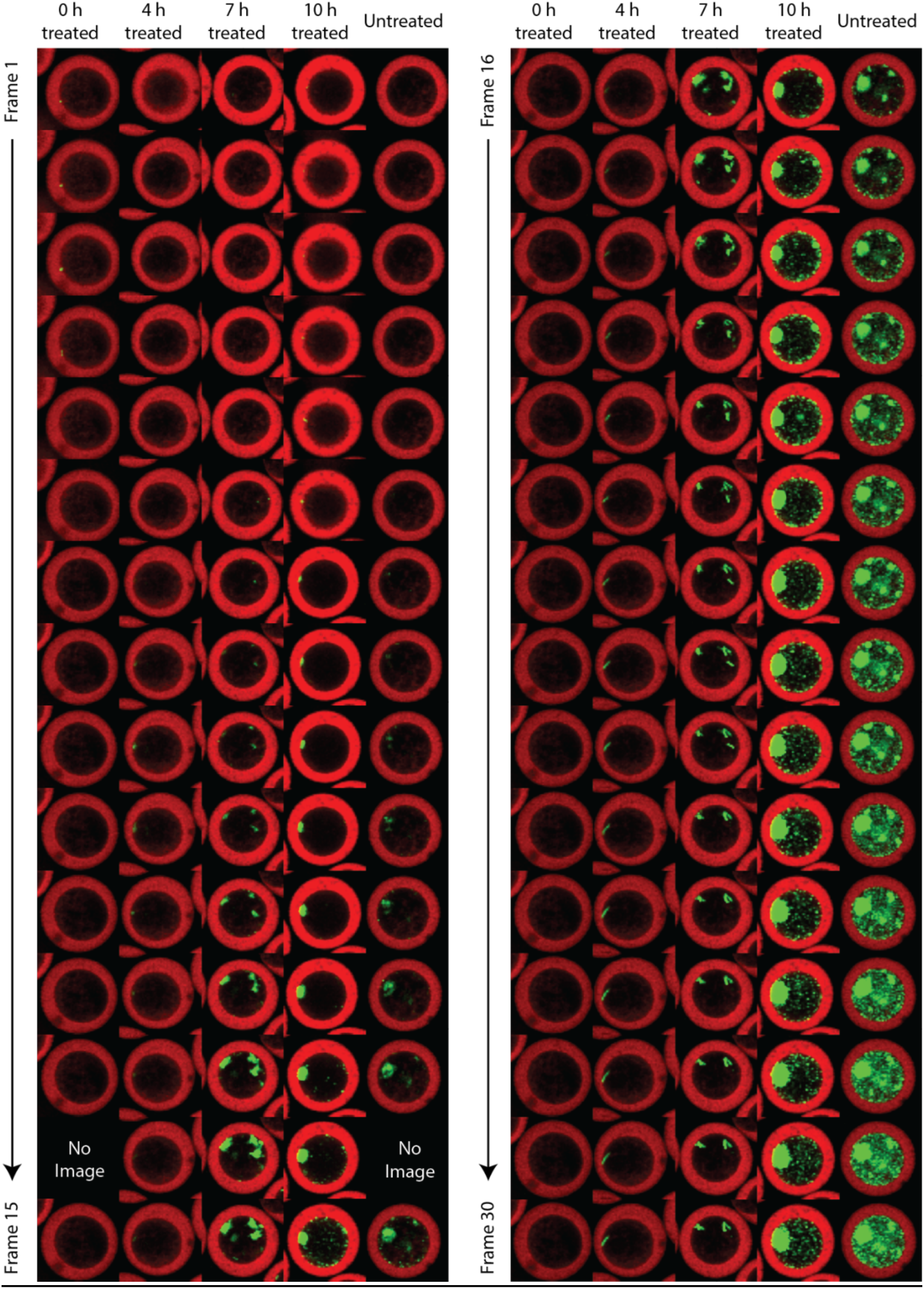
Representative capsule from each antibiotic treatment schedule shown for each individual imaging frame. Images represent a single slice taken at the capsule middle-plane via iCLSM. Capsule shells are shown in red (Cy5.5) and *P. aeruginosa* is in green (GFP). Capsule diameters: approx. 80 µm.

**Supplementary Figure 6:**
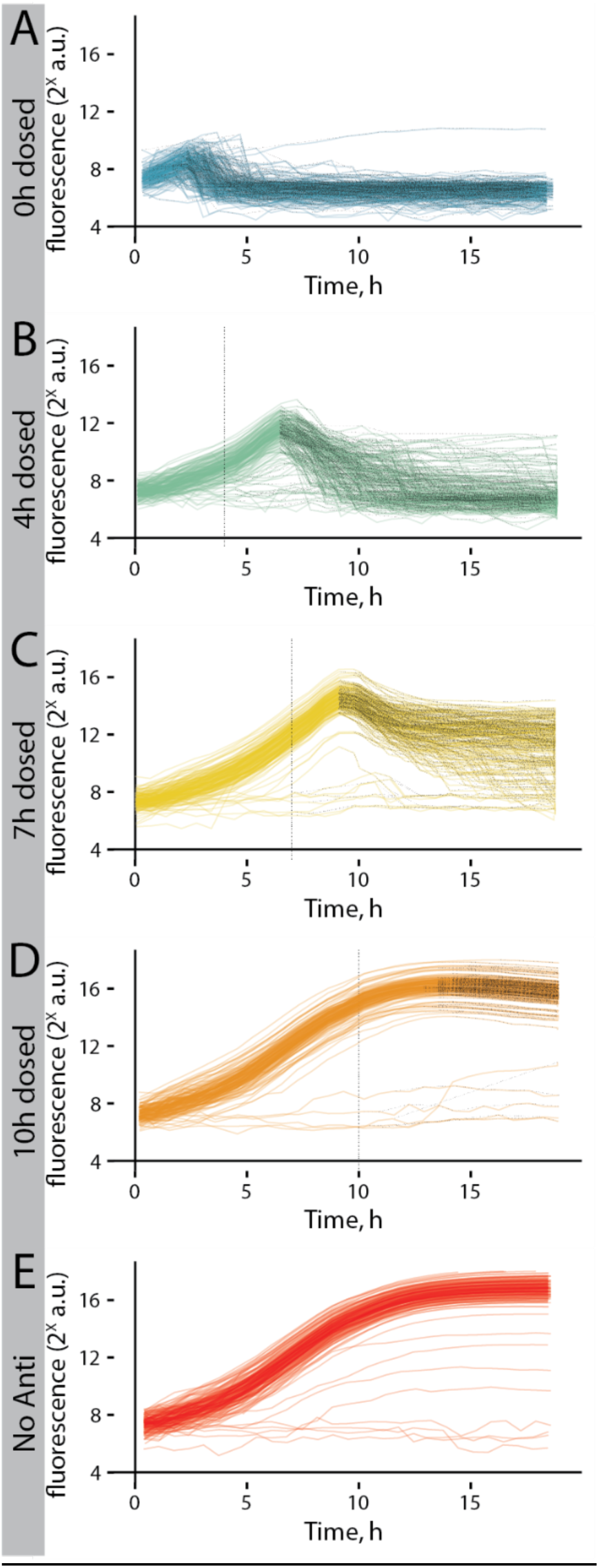
All individual capsule growth curves from each antibiotic treatment schedule tested. Black dashed lines overlay portions of each curve with a negative slope, representing decreased fluorescence.

## Notes

### Competing Interest Statement

The authors have declared no competing interest.

## REFERENCES CITED

1 Lyczak, J. B., Cannon, C. L. & Pier, G. B. Lung infections associated with cystic fibrosis. Clinical microbiology reviews 15, 194–222 (2002).

2 Malhotra, S., Hayes Jr, D. & Wozniak, D. J. Cystic fibrosis and Pseudomonas aeruginosa: the host-microbe interface. Clinical microbiology reviews 32, 10.1128/cmr. 00138-00118 (2019).

3 Davis, K. M. & Isberg, R. R. Defining heterogeneity within bacterial populations via single cell approaches. Bioessays 38, 782–790 (2016).

4 Amann, R. & Fuchs, B. M. Single-cell identification in microbial communities by improved fluorescence in situ hybridization techniques. Nature Reviews Microbiology 6, 339–348 (2008).

5 Moffitt, J. R. et al. High-throughput single-cell gene-expression profiling with multiplexed error-robust fluorescence in situ hybridization. Proceedings of the National Academy of Sciences 113, 11046–11051 (2016).

6 Beatty, K. E., Xie, F., Wang, Q. & Tirrell, D. A. Selective dye-labeling of newly synthesized proteins in bacterial cells. J Am Chem Soc 127, 14150–14151 (2005). 10.1021/ja054643w

7 Basu, S., Campbell, H. M., Dittel, B. N. & Ray, A. Purification of specific cell population by fluorescence activated cell sorting (FACS). Journal of visualized experiments: JoVE (2010).

8 Song, Y., Yin, H. & Huang, W. E. Raman activated cell sorting. Current opinion in chemical biology 33, 1–8 (2016).

9 Kashima, Y. et al. Single-cell sequencing techniques from individual to multiomics analyses. Experimental & Molecular Medicine 52, 1419–1427 (2020).

10 Hartmann, R. et al. Quantitative image analysis of microbial communities with BiofilmQ. Nature microbiology 6, 151–156 (2021).

11 Vidakovic, L., Singh, P. K., Hartmann, R., Nadell, C. D. & Drescher, K. Dynamic biofilm architecture confers individual and collective mechanisms of viral protection. Nature microbiology 3, 26–31 (2018).

12 Connell, J. L. et al. Probing prokaryotic social behaviors with bacterial “lobster traps”. MBio 1, e00202–00210 (2010).

13 Wessel, A. K., Hmelo, L., Parsek, M. R. & Whiteley, M. Going local: technologies for exploring bacterial microenvironments. Nature Reviews Microbiology 11, 337–348 (2013).

14 Akiyama, T. et al. Resuscitation of Pseudomonas aeruginosa from dormancy requires hibernation promoting factor (PA4463) for ribosome preservation. Proceedings of the National Academy of Sciences 114, 3204–3209 (2017).

15 Pratt, S. L. et al. DropSOAC: stabilizing microfluidic drops for time-lapse quantification of single-cell bacterial physiology. Frontiers in Microbiology 10, 2112 (2019).

16 Sart, S., Ronteix, G., Jain, S., Amselem, G. & Baroud, C. N. Cell culture in microfluidic droplets. Chemical Reviews 122, 7061–7096 (2022).

17 Anggraini, D. et al. Recent advances in microfluidic devices for single-cell cultivation: methods and applications. Lab on a Chip (2022).

18 Tao, Y. et al. Rapid, targeted and culture-free viral infectivity assay in drop-based microfluidics. Lab on a Chip 15, 3934–3940 (2015).

19 Rotem, A. et al. High-throughput single-cell labeling (Hi-SCL) for RNA-Seq using drop-based microfluidics. PloS one 10, e0116328 (2015).

20 Leonaviciene, G., Leonavicius, K., Meskys, R. & Mazutis, L. Multi-step processing of single cells using semi-permeable capsules. Lab on a Chip 20, 4052–4062 (2020).

21 van Zee, M. et al. High-throughput selection of cells based on accumulated growth and division using PicoShell particles. Proceedings of the National Academy of Sciences 119, e2109430119 (2022).

22 Jansen, L. E., Negrón-Piñeiro, L. J., Galarza, S. & Peyton, S. R. Control of thiol-maleimide reaction kinetics in PEG hydrogel networks. Acta biomaterialia 70, 120-128 (2018).

23 Nair, D. P. et al. The thiol-Michael addition click reaction: a powerful and widely used tool in materials chemistry. Chemistry of Materials 26, 724–744 (2014).

24 Kim, J. et al. Characterization of the crosslinking kinetics of multi-arm poly (ethylene glycol) hydrogels formed via Michael-type addition. Soft matter 12, 2076–2085 (2016).

25 Siltanen, C. et al. One step fabrication of hydrogel microcapsules with hollow core for assembly and cultivation of hepatocyte spheroids. Acta biomaterialia 50, 428–436 (2017).

26 Fattahi, P. et al. Core–shell hydrogel microcapsules enable formation of human pluripotent stem cell spheroids and their cultivation in a stirred bioreactor. Scientific reports 11, 1–13 (2021).

27 Rotem, A., Abate, A. R., Utada, A. S., Van Steijn, V. & Weitz, D. A. Drop formation in non-planar microfluidic devices. Lab on a Chip 12, 4263–4268 (2012).

28 Schindelin, J., et al. Fiji: an open-source platform for biological-image analysis. Nature methods 9, 676-682 (2012).

29 Pistori, H., Pistori, J. & Costa, E. R. in Workshop Software Livre, Porto Alegre. Anais do 6o Workshop de Software Livre.

30 Ferreira, L. A. & Teixeira, J. A. Salt effect on the aqueous two-phase system PEG 8000− Sodium sulfate. Journal of Chemical & Engineering Data 56, 133–137 (2011).

31 Nivens, D. E., Ohman, D. E., Williams, J. & Franklin, M. J. Role of alginate and its O acetylation in formation of Pseudomonas aeruginosa microcolonies and biofilms. Journal of bacteriology 183, 1047–1057 (2001).

32 Weinstein, M. P. Methods for dilution antimicrobial susceptibility tests for bacteria that grow aerobically. (No Title) (2018).

33 Ershov, D. et al. TrackMate 7: integrating state-of-the-art segmentation algorithms into tracking pipelines. Nature methods 19, 829–832 (2022).

34 Allaire, J. RStudio: integrated development environment for R. *Boston*, MA 770, 165–171 (2012).

35 Ihaka, R. & Gentleman, R. R: a language for data analysis and graphics. Journal of computational and graphical statistics 5, 299–314 (1996).

36 Wickham, H. & Wickham, M. H. Package tidyverse. Easily Install and Load the ‘Tidyverse (2017).

37 Wickham, H., Chang, W. & Wickham, M. H. Package ‘ggplot2’. Create elegant data visualisations using the grammar of graphics. Version 2, 1–189 (2016).

38 Jockers, M. L., Thalken, R., L Jockers, M. & Thalken, R. Introduction to dplyr. Text Analysis with R: For Students of Literature, 121–132 (2020).

39 Robinson, D., et al. Package ‘broom’. (2015).

40 Sánchez-Tapia, A. et al. in High Performance Computing: 4th Latin American Conference, CARLA 2017, Buenos Aires, Argentina, and Colonia del Sacramento, Uruguay, September 20-22, 2017, Revised Selected Papers 4. 218–232 (Springer).

41 Wickham, H., Wickham, M. H. & RColorBrewer, I. Package ‘scales’. R package version 1 (2016).

42 Bates, D., et al. Package ‘lme4’. *URL* http://lme4*. r-forge. r-project. org* (2009).

43 Kuznetsova, A., Brockhoff, P. B. & Christensen, R. H. B. Package ‘lmertest’. R package version 2, 734 (2015).

44 Cheng, Q. et al. Tunable Janus geometric morphology from aqueous two-phase systems on a superhydrophobic substrate. Journal of Materials Chemistry A 11, 4155–4161 (2023).

45 Headen, D. M., Aubry, G., Lu, H. & García, A. J. Microfluidic-based generation of size-controlled, biofunctionalized synthetic polymer microgels for cell encapsulation. Advanced materials 26, 3003–3008 (2014).

46 Agarwal, P. et al. One-step microfluidic generation of pre-hatching embryo-like core–shell microcapsules for miniaturized 3D culture of pluripotent stem cells. Lab on a Chip 13, 4525–4533 (2013).

47 Wang, X. et al. Proliferation and differentiation of mouse embryonic stem cells in APA microcapsule: A model for studying the interaction between stem cells and their niche. Biotechnology progress 22, 791–800 (2006).

48 Zhang, W. et al. A novel core–shell microcapsule for encapsulation and 3D culture of embryonic stem cells. Journal of materials chemistry B 1, 1002–1009 (2013).

49 Watanabe, T., Motohiro, I. & Ono, T. Microfluidic formation of hydrogel microcapsules with a single aqueous core by spontaneous cross-linking in aqueous two-phase system droplets. Langmuir 35, 2358–2367 (2019).

50 DeLeon, S. et al. Synergistic interactions of Pseudomonas aeruginosa and Staphylococcus aureus in an in vitro wound model. Infection and immunity 82, 4718–4728 (2014).

51 Phelps, E. A. et al. Maleimide cross-linked bioactive peg hydrogel exhibits improved reaction kinetics and cross-linking for cell encapsulation and in situ delivery. Advanced materials 24, 64–70 (2012).

52 Tang, T.-C. et al. Hydrogel-based biocontainment of bacteria for continuous sensing and computation. Nature chemical biology 17, 724–731 (2021).

53 Li, Y. et al. Construction of multilayer alginate hydrogel beads for oral delivery of probiotics cells. International journal of biological macromolecules 105, 924–930 (2017).

54 Quinn, R. A. et al. A Winogradsky-based culture system shows an association between microbial fermentation and cystic fibrosis exacerbation. The ISME journal 9, 1024–1038 (2015).

55 Patini, R. et al. The effect of different antibiotic regimens on bacterial resistance: a systematic review. Antibiotics 9, 22 (2020).

56 Wicha, S. G. et al. From therapeutic drug monitoring to model-informed precision dosing for antibiotics. Clinical Pharmacology & Therapeutics 109, 928–941 (2021).

57 Krismer, B. et al. Nutrient limitation governs Staphylococcus aureus metabolism and niche adaptation in the human nose. PLoS pathogens 10, e1003862 (2014).

58 Lewis, K. Persister cells. Annual review of microbiology 64, 357–372 (2010).

